# A Cooperative Mechanism of Eukaryotic Transcription Factor Target Search

**DOI:** 10.64898/2026.03.09.709760

**Authors:** Joseph V.W. Meeussen, Wim Pomp, Wim J. de Jonge, Davide Mazza, Tineke L. Lenstra

**Affiliations:** Division of Gene Regulation, The Netherlands Cancer Institute, Oncode Institute, Amsterdam, The Netherlands; Università Vita-Salute San Raffaele, IRCCS Ospedale San Raffaele, Experimental Imaging Center, Milan, Italy; Division of Single Cell Genomics, Princes Maxima Centrum; Utrecht, the Netherlands

**Keywords:** Gene Regulation, Transcription, Transcription Factor, Target Search, Single-molecule Microscopy, Dynamics

## Abstract

Rapid gene activation requires transcription factors (TFs) to locate their target motifs within vast genomes. In bacteria, TF target search is accelerated by combining 3D diffusion with 1D sliding, called facilitated diffusion, yet whether eukaryotic TFs rely on similar strategies has remained unresolved due to the lack of direct measurements in living cells. Here, we directly visualize eukaryotic TF target search in living cells by labeling a single TF per nucleus and visualizing its binding to its endogenous locus. Using the budding yeast TF Gal4, we find that efficient target localization occurs near the diffusion limit and does not require facilitated diffusion. Instead, rapid association requires cooperative self-interactions mediated by an intrinsically-disordered central region (IDR), independent of the activation domain. Replacing the Gal4 IDR with human self-interacting IDRs (EWS or FUS) restores efficient search, demonstrating that self-interactions are a general and portable feature for search. A second structured dimerization domain further cooperatives with the IDR to stabilize binding at neighboring motifs, revealing two mechanistically separable forms of cooperativity that together govern TF function. These findings highlight that facilitated diffusion is not strictly required and establish cooperative IDR-driven self-interactions as a key mechanism to enable rapid target recognition in eukaryotic cells.

## INTRODUCTION

Sequence-specific transcription factors (TFs) are essential for proper gene regulation by controlling transcription of their target genes.^1,2^ To enable fast and dynamic transcriptional responses, TFs must efficiently navigate through the crowded nuclear environment and scan millions of non-specific DNA basepairs for the correct motif.^1,3,4^ At target sequences, TFs bind specifically to DNA with their DNA-binding domain (DBD). Subsequent interactions of the TF’s activation domain (AD) with other transcriptional proteins, including coactivators and members of the pre-initiation complex (PIC), allows for initiation of transcription by RNA polymerase II (RNA Pol II). Eventually, the TF releases from the DNA and diffuses back into the nucleoplasm, ready to find its next target.

The TF search process has been best characterized in bacteria. Already in the 1970’s, it was observed that the *lac* repressor (LacI) associates with its binding sites much faster than would be possible by random three-dimensional (3D) diffusion.^5^ This accelerated target binding can be achieved by facilitated diffusion, where 3D diffusion is interspersed with 1D sliding along the DNA.^6–8^ Live-cell single-molecule imaging showed that LacI spends ∼90% of its time sliding along DNA, resulting in a search time of 3-5 min to find one target.^7,8^

However, whether eukaryotes use similar or fundamentally different search strategies to those in bacteria remains unclear. It has often been proposed that eukaryotic TFs also exploit facilitated diffusion, because the predicted TF search time by 3D diffusion only in large mammalian nuclei is in the order of days.^1^ While 1D sliding has been observed *in vitro*,^9,10^ it has not been directly demonstrated *in vivo*. Moreover, intrinsically disordered regions (IDRs) have recently been implicated in promoter selectivity, leading to the speculation that IDRs could regulate target recognition,^11–14^ but direct evidence that such IDRs drive target search in living eukaryotic cells is still lacking. Other potential search mechanisms include compact or guided exploration, engaging in homo- or heterotypic interactions, regulation of DNA accessibility by chromatin compaction, or nuclear partitioning.^1,4,15–19^ Alternatively, eukaryotic cells may simply express many TF copies to achieve rapid target binding. However, in the absence of any direct search time measurement, the search strategies of eukaryotic TFs remain poorly understood.

Single-molecule tracking (SMT) has proven to be a powerful method to study the *in vivo* dynamics of single TFs. However, interpretation of the observed TF kinetics is challenging. As SMT experiments lack information about the genomic context of the observed TF DNA-binding events, they comprise a mix of non-specific and specific interactions at sites with different DNA sequences.^20,21^ Moreover, since the number of labeled TF molecules and target sites is unknown, the TF search times can only be inferred using various assumptions.^22^ Additionally, the gene-agnostic nature of conventional SMT has prevented testing of any hopping or sliding mechanisms *in vivo.* Here, we labeled *only one* TF molecule per nucleus and visualized the association and dissociation kinetics of this single TF molecule at a single endogenous target locus. This one-TF-one-target approach provides gene-specific residence times, and allowed us, for the first time, to directly measure and mechanistically study the *in vivo* search of a eukaryotic TF molecule for an endogenous target locus.

Tracking a single TF shows that the eukaryotic TF Gal4 finds its *GAL* target locus in budding yeast with an average search time of approximately 5 minutes. This search time is close to, but approximately 4 times slower than the 3D diffusion limit. Although target search is efficient, it does not require 1D sliding or 3D hopping. Moreover, the transcriptional activation domain and a putative second dimerization domain identified by Alphafold3 do not affect Gal4 search kinetics. Mechanistically, eukaryotic target search requires cooperative interactions by an intrinsically disordered region (IDR) outside the DNA binding, dimerization and activation domains. Replacement of the Gal4-IDR by IDRs from mammalian EWS1 and FUS rescues TF search, which points to a mechanism where eukaryotic TF search is driven by IDR-mediated self-interactions. Moreover, our results reveal that both TF DNA binding stability and TF target search are cooperative, but they are partly independent and regulated by distinct domains. We propose a mechanism where Gal4 search is facilitated by IDR-mediated recognition of other Gal4-bound molecules that are localized at the *GAL* locus, after which self-interactions by IDRs and oligomerization domains allow for stable binding to neighboring TFBSs. Our findings indicate that compared to the facilitated diffusion observed in prokaryotes, eukaryotic TFs employ a conceptually different target search mechanism which depends on IDR-mediated cooperativity.

## RESULTS

### Direct observation of TF search at an endogenous target locus in living cells

Building on prior work,^23^ we used the budding yeast TF Gal4 and its *GAL* target locus as a model system. Gal4 contains a DNA-binding domain (DBD), a dimerization domain, a ∼700 amino-acid central region with unknown function and an activation domain (Figure 1A). Endogenous Gal4 was C-terminally tagged with HaloTag-V5. Importantly, sparse labeling with the bright and photostable dye JFX650H together with post-analysis filtering ensured that *only one* Gal4-HaloTag-V5 molecule was labeled per nucleus (Figures S1A–S1C, Video S1, Methods).^24^ The target *GAL* locus contains 3 genes – *GAL1*, *GAL10* and *GAL7* – and harbors 6 out of 15 genomic Gal4 transcription factor binding sites (TFBSs).^25^ The *GAL* locus was labeled with 128x*tetO* tetR-ymScarletI downstream of *GAL1* (Figures 1B and S1D).^26,27^ It was kept in focus with our focus-feedback tracking algorithm,^23^ which uses an active feedback loop to refocus on the DNA label between subsequent acquisitions (Figure S1A). As previously,^23^ the DNA-binding dynamics of Gal4 at its *GAL* target locus were recorded for a period of 20 minutes per cell, using a 5 s interval between frames and a relatively long exposure time of 100 ms, blurring freely diffusing TF molecules. Intensity traces of Gal4 molecules detected within 0.4 µm (*xy*-distance) from the *GAL* DNA label were binarized with an intensity threshold to discriminate between the “unbound” and “bound” states (Figures 1C, 1D and S1E, Video S1, Methods). We previously showed that Gal4 binding event of multiple seconds are able to activate transcription, indicating they represent specific binding events to TFBS.^23^ We therefore defined Gal4 molecules as bound only if binding events lasted ≥2 frames (≥5 s) to limit contributions of non-specific binding, clustered Gal4 molecules and transient associations (Methods). Together, these unique conditions ensure that each trace captures the binding of an individual TF molecule to the locus. The durations of the bound and unbound states are direct measurements of the respective residence times and target search times of Gal4 at the *GAL* locus.

**Figure 1.**
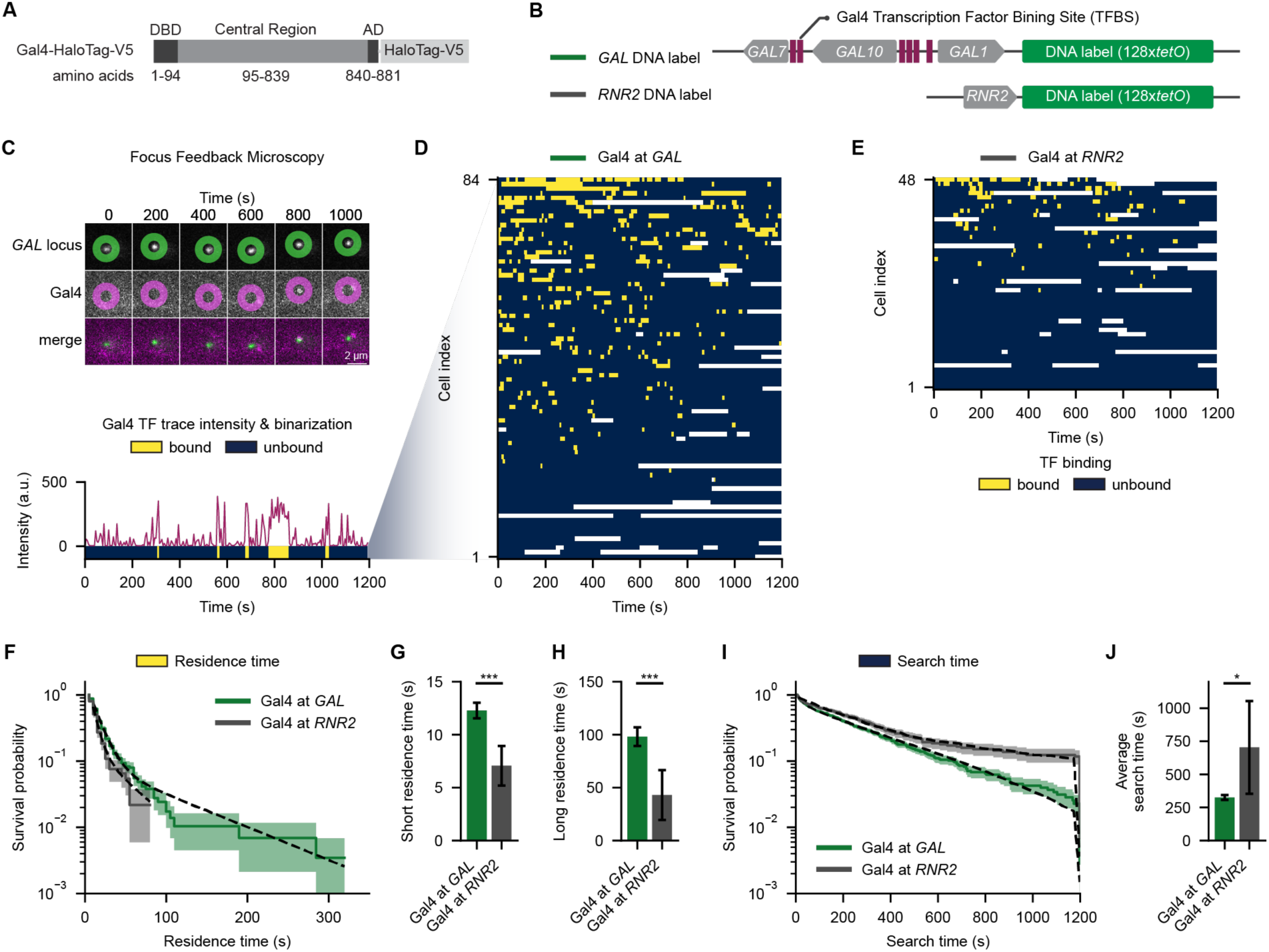
Direct observation of Gal4 residence times and target search times at a single endogenous locus in living cells. (A) Schematic of the transcription factor (TF) Gal4-HaloTag-V5, which is conjugated with JFX650H. (B) Schematic of the *GAL* target locus (top), containing six Gal4 transcription factor binding sites (TFBS; purple bars), and the *RNR2* locus (bottom), without any Gal4 binding sites. Live-cell visualization of these loci is established by genomic integration of 128x*tetO* repeats, forming a fluorescent DNA label together with tetR-ymScarletI (DNA label; green). (C) Direct observation of a single Gal4 molecule binding at the *GAL* locus (top), with raw TF intensity trace (bottom; purple) and binarization (bottom; blue = unbound, yellow = bound). (D) Heatmap of binarized intensity traces capturing target search and residence time durations of single Gal4 molecules at the *GAL* locus (blue = unbound; yellow = bound; white = no data). (E) Same as (D) for the *RNR2* locus. (F) Survival probability distributions of residence times of Gal4 at the *GAL* locus (green; *n*=84 traces) and at the *RNR2* locus (grey; *n*=48 traces). Solid lines indicate raw durations, shaded areas represent bootstrapped 95% confidence intervals and dotted lines show the biexponential fits. (G, H) Residence time fit parameters of the survival probability distributions in (F), indicating (G) short residence time, (H) long residence time. Error bars represent 95% confidence interval from the fit. (I) Same as (F) for the Gal4 search times. (J) Average search times from the fits of the survival probability distributions in (I). Error bars represent 95% confidence interval from the fit.

The survival probability distributions of the residence times were best fit using a bi-exponential model, indicating two binding populations (Figure 1F, Methods). Similar to our previous measurements,^23^ many Gal4 molecules displayed short residence times (12.3 ± 0.7 s, mean ± 95% confidence interval), and a smaller fraction (0.07 ± 0.01) bound the *GAL* locus for prolonged periods (98.1 ± 8.9 s) (Figures 1G and 1H). To investigate the specificity of these DNA-binding dynamics, we repeated the experiment in a yeast strain where the DNA label was placed next to *RNR2* locus, which is located on a different chromosome without any proximal Gal4 binding sites (Figures 1B and S1D). The durations of both the short and long residence times of Gal4 at the non-target locus *RNR2* were much shorter (Figures 1E–1H and S1F), confirming the specificity of the measurements. Compared to WT Gal4 at the *GAL* locus, the short and long residence times at *RNR2* were closer in time scales and more difficult to distinguish, similar to the mutants described below. We therefore used the two-component fits to compare the average residence times in the remainder of the manuscript (Methods).

Next, we exploited the single-TF-single-gene setup to directly determine the durations in between binding events, which reflects the search time of individual Gal4 molecules at the *GAL* and *RNR2* loci *in vivo.* Direct measurements of Gal4 search time similarly showed a biexponential distribution (Figure 1I). The biexponential fit was used to determine the average search time, which for Gal4 at the *GAL* locus was 325 ± 19 s (mean ± 95% confidence interval) (Figure 1J, Methods). On average, a single Gal4 TF thus revisits the *GAL* locus within approximately 5 minutes after dissociating. As expected, at the non-target *RNR2*, Gal4 bound much less frequently (search time of 704 ± 350 s) (Figures 1I and 1J).

The observed *in vivo* Gal4 binding dynamics allow an estimation of the apparent binding affinity (*K_D_*) of Gal4 for the *GAL* locus of 11 ± 1 nM (mean ± 95% confidence interval, Methods), which is remarkably similar to the *in vitro* affinity of Gal4-DBD to cognate motifs of 3–35 nM.^28–31^ The similarity between these *in vivo* and *in vitro* affinities further validates our approach.

### Comparison of Gal4 search time with Smoluchowski-limit suggests Gal4 search is diffusion-limited

To understand whether facilitated diffusion is required for Gal4 to reach its target, we compared the measured search time with the Smoluchowski-limit, which describes the search time for a TF to reach its target by unhindered random 3D diffusion.^32^ To calculate this diffusion limit, we first measured the Gal4 diffusion rate using fast SMT with continuous exposure and 12 ms intervals, which revealed a diffusion coefficient of 0.31 ± 0.03 μm^2^/s (Figures S1G and S1H, Methods). Using this diffusion coefficient, the diffusion limit of a Gal4 dimer is ∼74 s (Methods). Although the measured Gal4 search time of 325 ± 19 s converges to the diffusion limit, it remains ∼4.4 times slower. Varying the binarization threshold, correcting for photobleaching, adjusting motif number, or inclusion of potentially non-specific binding events of 1 frame influences the exact fold change (Figures S7, S8, S9, Methods), but in all cases the search time of Gal4 is diffusion-limited. The slower rather than faster search time suggest that, contrary to bacterial TFs, facilitated diffusion that includes a combination of 3D and 1D diffusion may not be required for the TF search process in eukaryotes. Nevertheless, given the numerous non-specific interactions and presence of other target motifs which may decelerate Gal4 search, the approximation of the diffusion limit suggest Gal4 search is very efficient.

### Gal4 search does not depend on sliding and hopping

To further understand the search mechanisms of Gal4 and to directly test the contribution of 1D sliding and 3D hopping, we designed several mutated promoter variants. The endogenous *GAL* locus contain six TFBSs, of which two in the *GAL7* and four in *GAL1-10* promoter (*GAL* wild type, Figure 2A). First, to test if Gal4 hops locally between neighboring TFBSs, we scrambled all neighboring TFBSs, leaving one in the *GAL7* promoter and one in the *GAL1-10* promoter intact (1+1TFBS, Figure 2A). Second, to test the contribution of long-range 3D jumping, we scrambled all TFBSs in the *GAL7* promoter and left only a single TFBS in the *GAL1-10* promoter (1TFBS, Figure 2A). Third, to test the contribution of 1D sliding, a roadblock (tetR TFBS *tetO*) was placed adjacent to this remaining binding site (1TFBS+*tetO*, Figure 2A), blocking sliding from one side. This latter roadblock was previously used to investigate 1D sliding in bacteria.^8^

**Figure 2.**
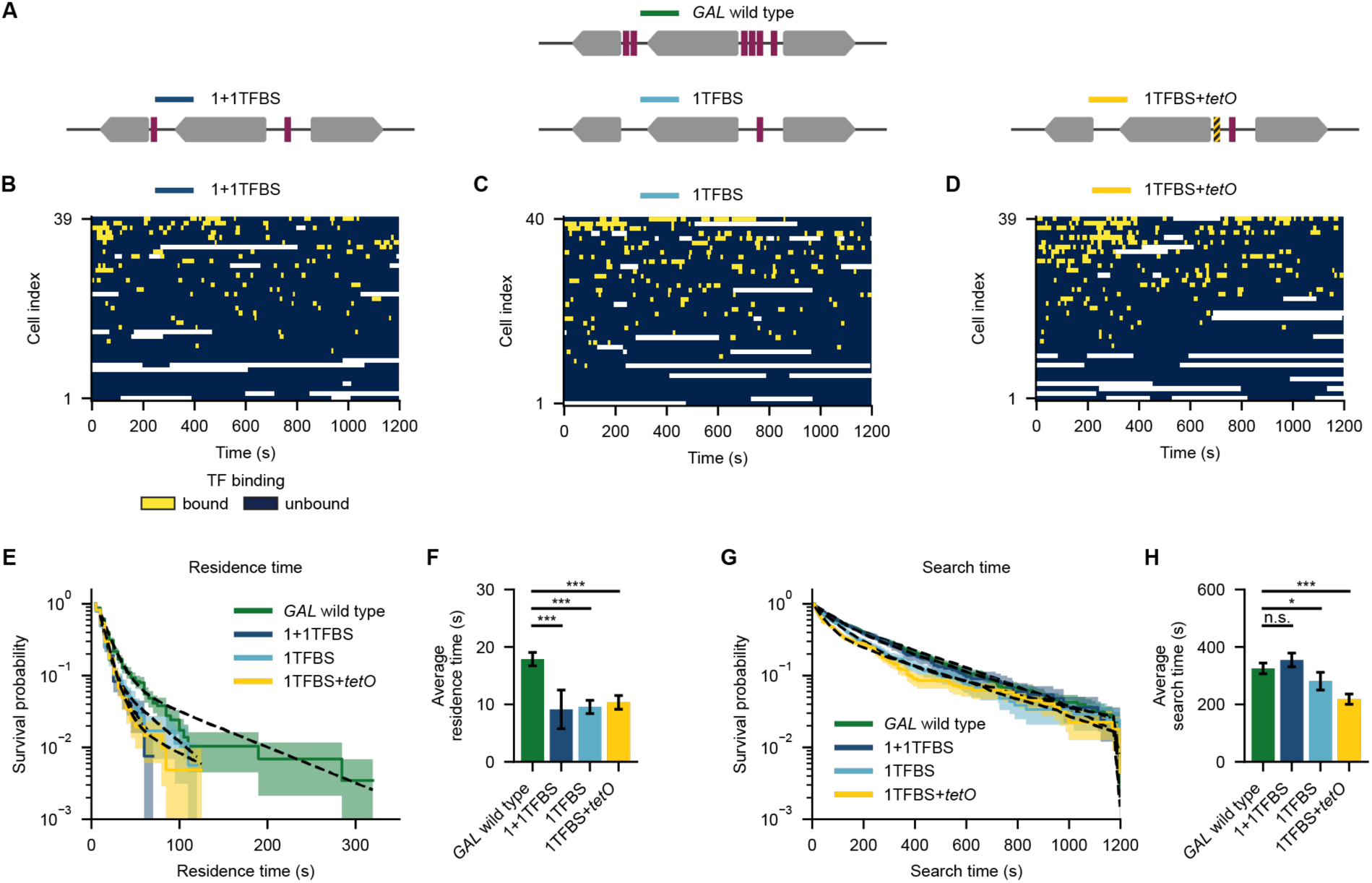
Gal4 search does not depend on sliding and hopping. (A) Schematic of *GAL* locus variants, with the *GAL* locus with the six endogenous Gal4 transcription factor binding sites (TFBS, purple bars) (*GAL* wild type, top), mutant with one in the *GAL7* promoter and one in *GAL1-10* promoter (1+1TFBS, bottom left), mutant with one TFBS in the *GAL1-10* promoter (1TFBS, bottom middle) and mutant with one TFBS in the *GAL1-10* promoter with a neighboring *tetO* sequence 19 bp upstream (1TFBS+*tetO*, bottom right). (B) Heatmap of binarized intensity traces capturing target search and residence time durations of single Gal4 molecules at the 1+1TFBS variant of the *GAL* locus (blue = unbound; yellow = bound; white = no data). (C, D) Same as (B) for the 1TFBS (C) or 1TFBS+*tetO* (D) variants of the *GAL* locus. (E) Survival probability distributions of residence times of Gal4 at *GAL* wild type locus (green; *n*=84 traces) and the variants 1+1TFBS (light blue; 39 traces), 1TFBS (dark blue; 40 traces) and 1TFBS+*tetO* (yellow; *n*=39 traces). Solid lines indicate raw durations, shaded areas represent bootstrapped 95% confidence intervals and dotted lines show the biexponential fits. (F) Average residence times from the fits of the survival probability distributions in (E). Error bars represent 95% confidence interval from the fit (G) Same as (E) for the search times. (H) Average search times from the fits of the survival probability distributions in (G). Error bars represent 95% confidence interval from the fit.

Traces of Gal4 binding at the 1+1TFBS, 1TFBS and 1TFBS+*tetO GAL* locus variants all showed approximately 50% shorter Gal4 binding events compared to those at the endogenous *GAL* locus (9.1 ± 3.4 s for 1+1TFBS, 9.6 ± 1.2 s for 1TFBS and 10.3 ± 1.2 s for 1TFBS+*tetO* vs 17.9 ± 1.2 s for *GAL* wild type) (Figures 2B–2F and S2A–S2C). Apart from further validating the gene-specificity of the approach, these reduced residence times indicate that neighboring binding sites in the wild type promoter support longer Gal4 residence times. These longer residence times may arise from rapid rebinding after release, which would be captured as a longer binding event. Alternatively, Gal4 DNA-binding may be stabilized by self-interactions with neighboring bound Gal4 dimers.

Surprisingly, even though the number of Gal4 binding sites was reduced from six to one or two, the search times for these promoter variants were very similar to those for the endogenous *GAL* locus (Figure 2G), with an average of search time of 355 ± 24 s for 1+1TFBS, 281 ± 31 s for 1TFBS and 218 ± 18 s for 1TFBS+*tetO* (Figure 2H). The absence of an increase in search time of having additional proximal or distal TFBSs or placing a neighboring *tetO* site shows that local hopping, long-range jumping, or 1D sliding contributes little to Gal4 search. This indicates that Gal4 uses alternative mechanisms than facilitated diffusion to find its target. Moreover, if these TFBSs would act independently, it would be expected that reducing the number of TFBSs would increase the search time for the locus. Importantly, the absence of this increase indicates that Gal4 recognizes a larger target size than the TFBSs itself.

### Stable DNA binding and efficient target search relies on cooperative interactions

We previously showed that Gal4 can self-interact,^27^ and DNA-bound Gal4 molecules could potentially be a larger target for recognition. In addition, Gal4 forms clusters which colocalize with the *GAL* locus.^27^ We therefore hypothesized that Gal4 recognizes either other DNA-bound or clustered Gal4 molecules at the locus through cooperative self-interactions.

To directly test how Gal4 cooperativity could contribute to Gal4 DNA-binding kinetics at the *GAL* locus, we mutated the *GAL4* promoter, which reduced its concentration to background levels (Figures 3A) and impeded growth on galactose (Figure S3A). Lowering the concentration of Gal4 molecules reduced the Gal4 residence times (10.1 ± 2.1 s vs 17.9 ± 1.2 s) (Figures 3B–3D and S3B) and increased the time that a single Gal4 molecule requires to find the *GAL* locus from 325 ± 19 s to 429 ± 53 s (Figures 3E and 3F). We conclude that both to TF target binding and target search require Gal4 cooperativity.

**Figure 3.**
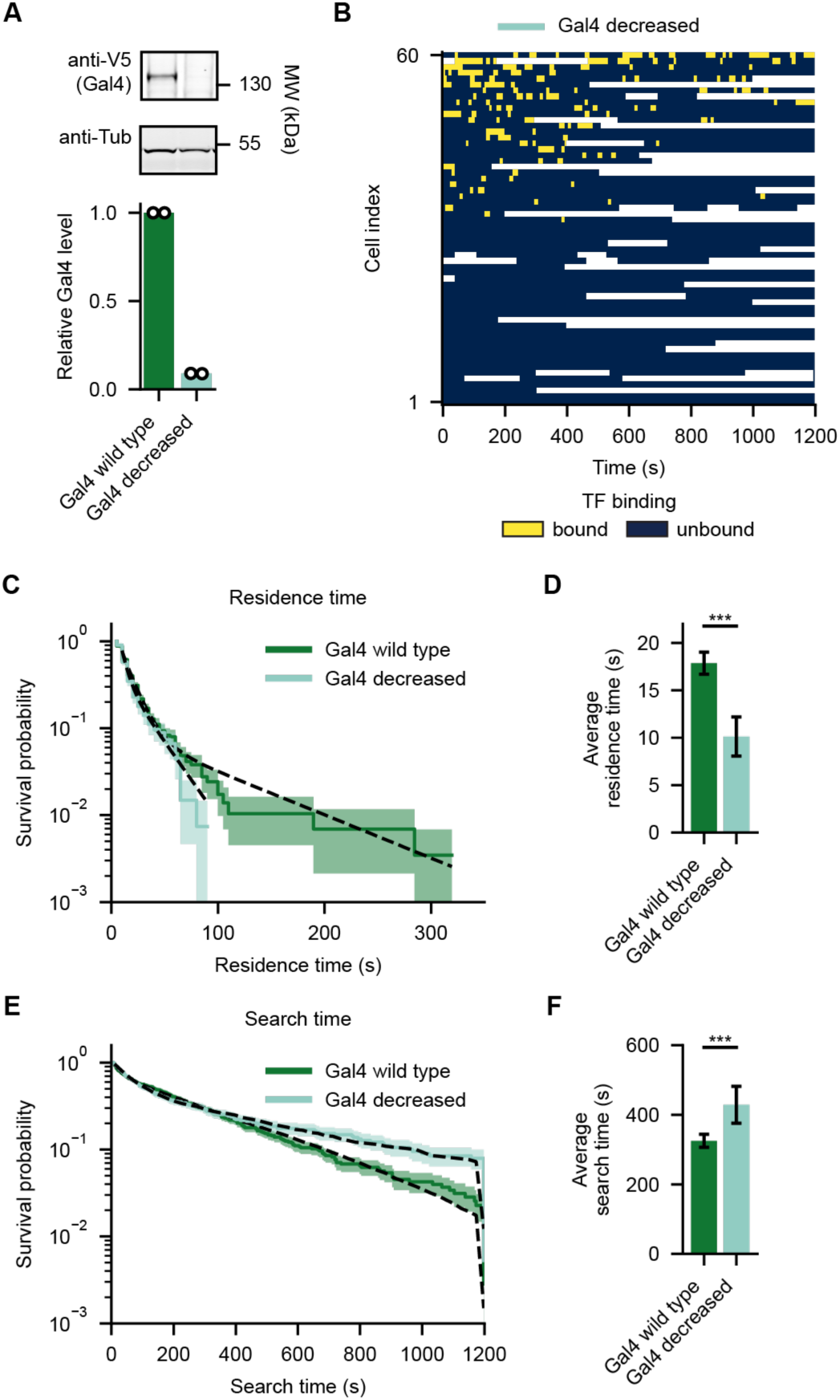
Gal4 search requires cooperative interactions. (A) Assessment of Gal4 protein levels by western blot of wild type *GAL4* promoter (Gal4 wild type, left) and mutated *GAL4* promoter (Gal4 decreased, middle) to verify the decreased Gal4 concentration upon mutating the *GAL4* promoter in the HaloTag-V5 background. Western blot image is a representative example of 2 independent experiments. The anti-V5 antibody signals are normalized to anti-Tubulin and to the expression of Gal4 wild type, resulting in the bar graph, in which open circles represent the results of individual replicate experiments, and the colored bars indicate their mean. (B) Heatmap of binarized intensity traces capturing target search and residence time durations of single Gal4 molecules at the *GAL* locus at a decreased Gal4 concentration (blue = unbound; yellow = bound; white = no data). (C) Survival probability distributions of residence times of Gal4 at the *GAL* locus at wild type (green; *n*=84 traces) and decreased (light green; *n*=60 traces) concentrations. Solid lines indicate raw durations, shaded areas represent bootstrapped 95% confidence intervals and dotted lines show the biexponential fits. (D) Average residence times from the fits of the survival probability distributions in (C). Error bars represent 95% confidence interval from the fit. (E) Same as (C) for the search times. (F) Average search times from the fits of the survival probability distributions in (E). Error bars represent 95% confidence interval from the fit.

### The Gal4 central region increases Gal4 residence times and decreases search time

To further understand how the observed Gal4 cooperativity facilitates stable DNA binding and efficient target search, we aimed to understand how the Gal4 DNA-binding kinetics are regulated by the different Gal4 domains. First, we compared full length Gal4 to the Gal4-DBD-only, which contains the dimerization domain, but lacks the central region and activation domain (Figure 4A). The DBD of Gal4 has often been used in combination with the viral activation domain VP16 to drive transgene expression in diverse organisms,^33,34^ highlighting the DBD is sufficient for function. The Gal4-DBD-only truncation mutant had protein levels comparable to full length Gal4 (Figure S4A) and was unable to grow on galactose (Figure S3A), as expected. Tracing this Gal4-DBD-only mutant at the *GAL* locus revealed predominantly short binding events and a decreased average residence time compared to wild-type (WT) Gal4 (7.9 ± 1.6 s vs 17.9 ± 1.2 s) (Figures 4B, 4D, 4E and S3B). In addition, analysis of the Gal4-DBD-only search time to the *GAL* genes revealed a large increase compared to WT (543 ± 105 s vs 325 ± 19 s) (Figures 4F and 4G). Although the DBD directly recognizes and interacts with the DNA motif, these results indicate that nonDBD regions are essential to increase the residence time and decrease the search time of Gal4 at the *GAL* target locus.

**Figure 4.**
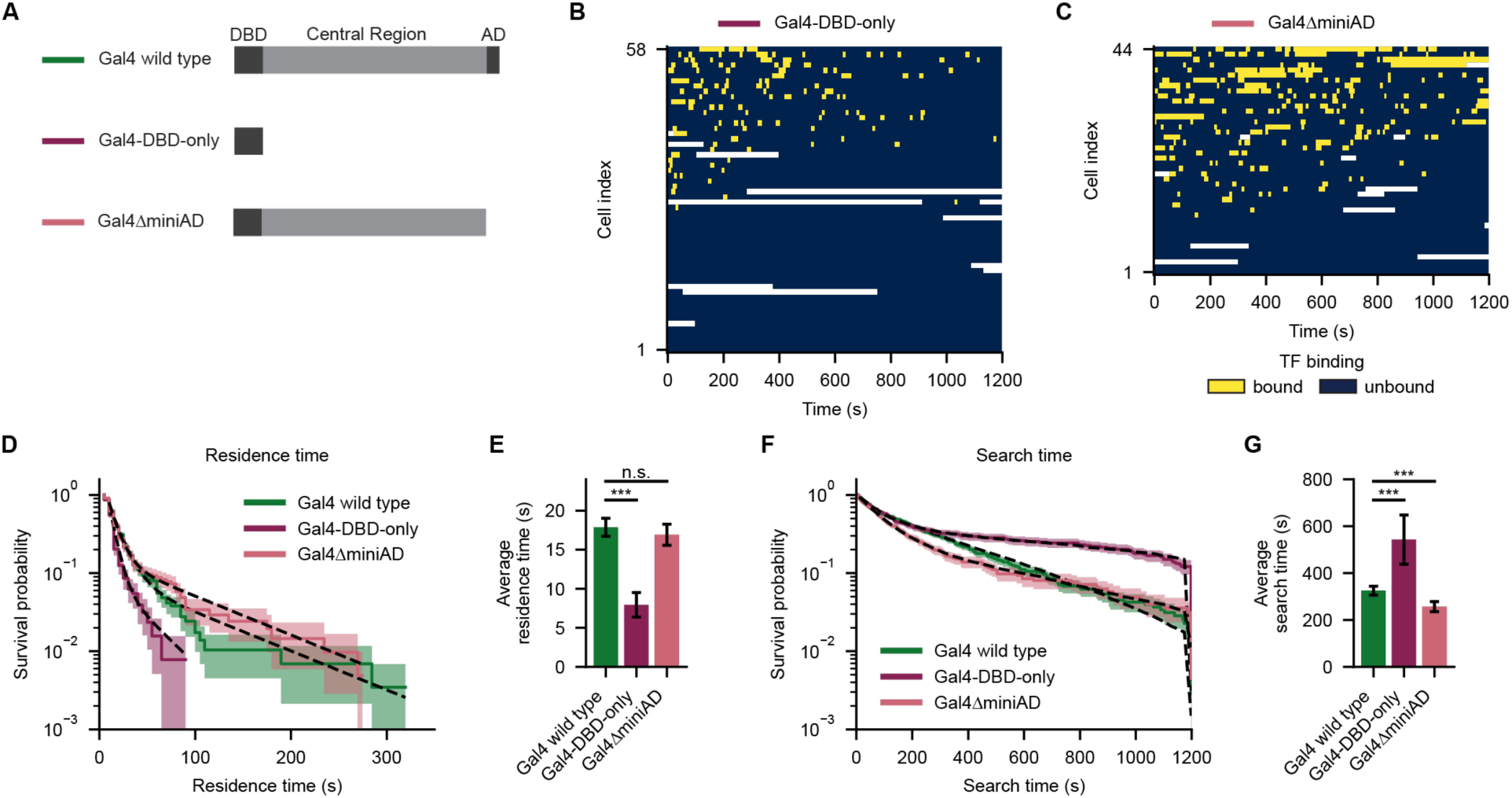
Gal4 requires regions outside its DNA-binding domain for prolonged target binding and efficient target search. (A) Schematic showing 2D-representations Gal4 wild type (top), Gal4-DBD-only (right top), and Gal4ΔminiAD (right bottom). (B) Heatmap of binarized intensity traces capturing target search and residence time durations of single Gal4-DBD-only molecules at the *GAL* locus (blue = unbound; yellow = bound; white = no data). (C) Same as (B) for Gal4ΔminiAD. (D) Survival probability distributions of residence times of Gal4 wild type (green; *n*=84 traces), Gal4-DBD-only (dark pink; *n*=58 traces) and Gal4ΔminiAD (pink; *n*=44 traces) at the *GAL* locus. Solid lines indicate raw durations, shaded areas represent bootstrapped 95% confidence intervals and dotted lines show the biexponential fits. (E) Average residence times from the fits of the survival probability distributions in (D). Error bars represent 95% confidence interval from the fit. (F) Same as (D) for the search times. (G) Average search times from the fits of the survival probability distributions in (F). Error bars represent 95% confidence interval from the fit.

Because these nonDBD regions include the activation domain, we asked whether either transcription activation itself or interactions of the activation domain (AD) with the transcriptional machinery affect the Gal4 DNA-binding dynamics. These AD-mediated interactions might result in destabilization of TF-DNA binding due to the action of chromatin remodelers or due to the accumulation of DNA supercoils as a result of RNA Pol II transcription.^31,35^ Alternatively, the TF AD might stabilize TF-DNA binding, for example by facilitating rapid rebinding after dissociation if the AD remains bound to coactivators such as Mediator.^36^

To perturb both transcriptional activation and cofactor interactions, we truncated a small part of the C-terminal activation domain (miniAD) of Gal4 (Figure 4A), which is essential for both transcriptional activation and Mediator interactions.^37^ Protein levels of the Gal4ΔminiAD were increased ∼2-fold compared to WT Gal4 (Figure S4A), and its inability to support growth on galactose was confirmed (Figure S3A). The *GAL* DNA binding kinetics of the Gal4ΔminiAD closely resembled those of full length Gal4, with similar average residence times (16.9 ± 1.3 s vs 17.9 ± 1.2 s) and search times (257 ± 21 vs 325 ± 19 s) (Figures 4B–4G and S4C), although minor deviations were observed in the shape of the search time distribution.

The effects on the DNA binding dynamics of these DBD-only and ΔminiAD results were validated with conventional SMT (30 ms exposure, 200ms interval, Methods). In agreement with the experiments at the *GAL* locus, the Gal4-DBD-only mutant substantially reduced the fraction of stable binding events throughout the nucleus compared to WT Gal4 (from 0.104 ± 0.025 to 0.045 ± 0.008), whereas the Gal4ΔminiAD did not majorly affect the stable binding fraction (from 0.104 ± 0.025 to 0.074 ± 0.023) (Figure S4D). These global SMT experiments also revealed that the fraction of transient non-specific interactions (∼1 s) of WT Gal4 was reduced for both the Gal4-DBD-only and Gal4ΔminiAD mutants (from 0.159 ± 0.040 in WT Gal4 to 0.029 ± 0.006 in Gal4-DBD-only and 0.101 ± 0.013 in Gal4ΔminiAD, Figure S4D). The C-terminal part of the Gal4 activation domain thus supports non-specific chromatin interactions, but this does not substantially affect its specific binding at the *GAL* locus. Moreover, since the stable DNA binding kinetics globally or at the *GAL* locus are largely unchanged in the Gal4ΔminiAD, we conclude that there is little feedback from the transcriptional machinery, either directly via cofactor interactions or indirectly via supercoiling mechanisms,^35^ on Gal4 association or dissociation. Lastly, the much larger kinetic effects of the Gal4-DBD-only compared to the Gal4ΔminiAD indicates that the central region has a key role in enhancing both the DNA-binding stability and the search efficiency of Gal4.

### Distinct domains of the central region regulate Gal4 DNA binding stability and search time

The importance of the Gal4 central region for the TF DNA binding kinetics warranted further dissection of this region. Although the Gal4-DBD with its dimerization domain (amino acids 1-100) has been crystalized, full-length Gal4 is notoriously difficult to purify, which has limited structural analysis of the full-length protein.^30,38^ Disorder prediction tools indicated that the C-terminal part of the central region is likely disordered, but that the N-terminal central region is structured.^27^ To obtain insight into the possible structure of the Gal4 central region, we applied Alphafold3 modeling on two full length Gal4 molecules in combination with DNA (Figures 5A and S5A).^39^ The resulting Alphafold3 model showed DNA binding by the Zn_2_Cys_6_ binuclear clusters of the Gal4-DBD (residues 7-40), which were bridged by the previously identified adjacent dimerization domain (residues 50-94), in line with the crystal structure.^30^ In agreement with the IDR-prediction tools,^27^ the C-terminal part of the central region and the activation domain were predicted to be mostly disordered (residues 651-881). In addition, the model also predicted a second dimerization domain in the central region (residues 95-650) with high confidence. AlphaBridge confidence scores for this second multimerization interface in the central region were similar to the confidence scores of the previously crystalized dimerization domain adjacent to the DBD (Figure S5B),^40^ suggesting bona fide interactions.^41^ Overall, these structure predictions revealed that the Gal4 central region contains a second structured di- or oligomerization domain and an IDR.

**Figure 5.**
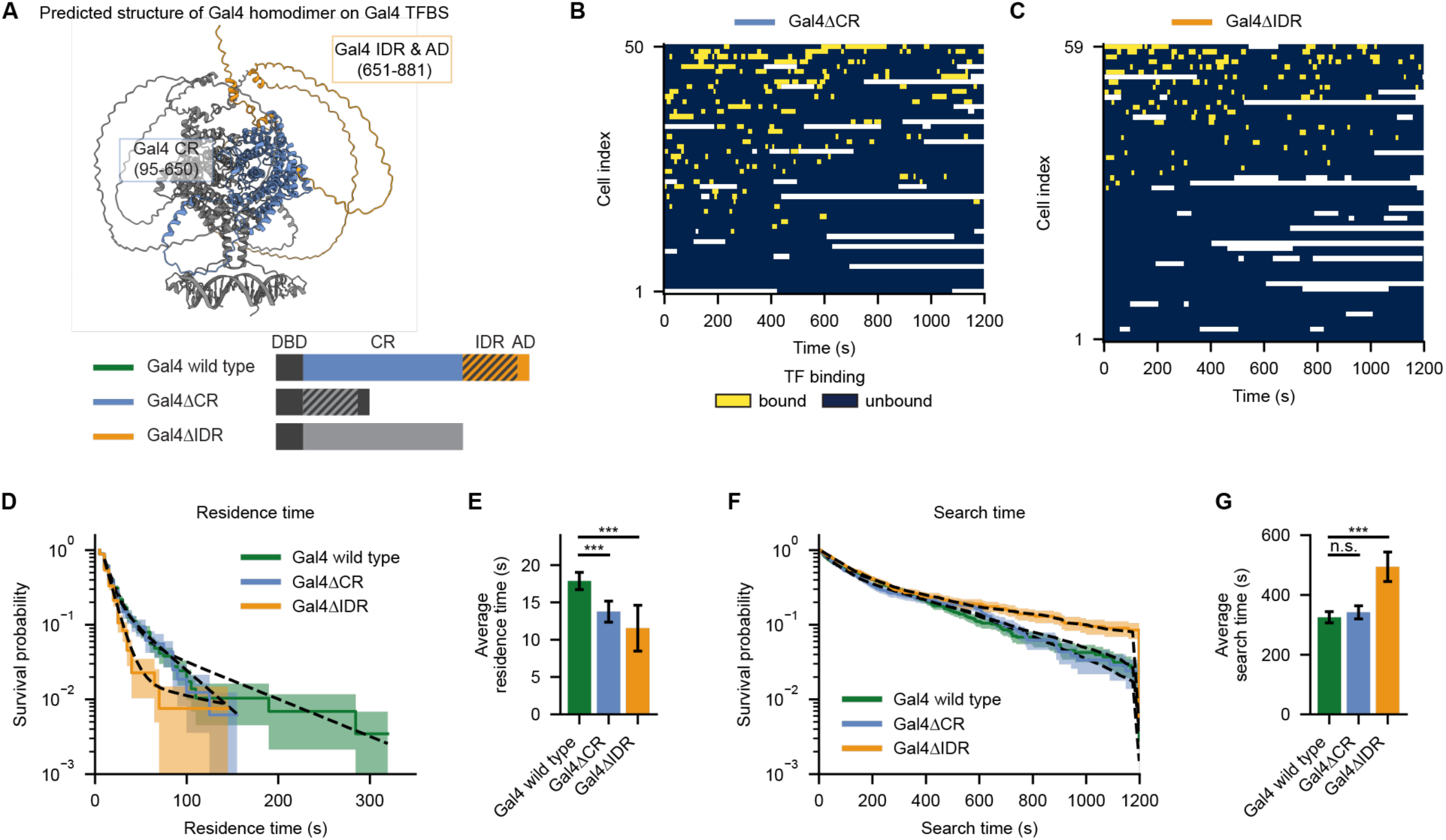
Efficient target search is mediated by the Gal4 IDR. (A) Top: Alphafold3 model of two full-length Gal4 molecules with DNA (sense + antisense sequence of TFBS 3 in the *GAL1-10* promoter) and 4 Zn^2+^ ions. The color of the structure highlights the Gal4 central region (CR; purple) and intrinsic disordered region and activation domain (IDR & AD; orange) including the amino acids corresponding to those regions. Bottom: 2D-representations of Gal4 wild type (top), Gal4ΔCR (middle) and Gal4ΔIDR (bottom). (B) Heatmap of binarized intensity traces capturing target search and residence time durations of single Gal4ΔCR molecules at the *GAL* locus (blue = unbound; yellow = bound; white = no data). (C) Same as (B) for Gal4ΔIDR. (D) Survival probability distributions of residence times of Gal4 wild type (green; *n*=84 traces), Gal4ΔCR (purple; *n*=50 traces) and Gal4ΔIDR (orange; *n*=59 traces) at the *GAL* locus. Solid lines indicate raw durations, shaded areas represent bootstrapped 95% confidence intervals and dotted lines show the biexponential fits. (E) Average residence times from the fits of the survival probability distributions in (D). Error bars represent 95% confidence interval from the fit. (F) Same as (D) for the search times. (G) Average search times from the fits of the survival probability distributions in (F). Error bars represent 95% confidence interval from the fit.

To understand the role of both regions in the Gal4 DNA binding kinetics, we truncated either the predicted dimerization domain in the central region (Gal4ΔCR) or deleted the entire C-terminal Gal4 IDR (Gal4ΔIDR, Figure 5A). Both mutants were expressed at similar or higher levels than wild type Gal4 (Figure S5A), but were unable to support growth on galactose (Figure S3A). Single-molecule tracing of the Gal4ΔCR and Gal4ΔIDR mutants showed that both mutants displayed shorter DNA residence times at the *GAL* target locus than WT Gal4 (13.8 ± 1.4 s for Gal4ΔCR and 11.6 ± 3.1 s for Gal4ΔIDR vs 17.9 ± 1.2 s for WT Gal4) (Figures 5B–5E, S5D and S5E). Although the DBD and adjacent dimerization domain supports DNA binding, the novel CR-dimerization domain and the IDR are thus important for prolonged DNA binding.

These mutants, however, displayed different effect on Gal4 search. Removal of the Gal4-CR did not affect Gal4 search (342 ± 22 s), whereas removal of the Gal4-IDR resulted in longer search times than wild type (494 ± 50 s vs 325 ± 19 s) (Figures 5F and 5G). The Gal4ΔIDR search times closely resembled those of the DBD-only (Figure 4F), indicating that efficient Gal4 target search is mainly mediated by this C-terminal IDR.

These and previous perturbations reveal that cooperative DNA binding requires di-/multimerization domains as well as the IDR, stabilizing Gal4 dimers on neighboring binding sites, while cooperative search is mediated by the IDR. We therefor uncover two mechanistically different forms of TF cooperativity that are regulated by distinct TF domains, independent of the DBD and activation domains.

### Replacement of Gal4-IDR by human IDRs restores TF target search

Given the role of the Gal4-IDR in regulating TF association and dissociation kinetics, we aimed to understand the mechanism and generality further. Since Gal4 has the ability to form clusters, we hypothesized that Gal4 cooperativity is established through IDR-mediated self-interactions, which would support prolonged binding at neighboring sites and more efficient target recognition. To test this, we replaced the entire C-terminal Gal4-IDR, including its activation domain, with the exogenous IDRs of EWS1 or FUS, well-known known for their ability to self-interact.^42^

Both the Gal4ΔIDR-EWS and Gal4ΔIDR-FUS were expressed at wild type levels (Figure S6A), but neither IDR variant rescued growth on galactose plates (Figure S6B), indicating these IDRs are unable to substitute for the Gal4 transcription activation domain in activating transcription of the *GAL* genes.

Single-molecule imaging showed that replacement of Gal4 IDR with that of FUS fully restored the DNA residence times to wild type (17.0 ± 1.3 s vs 17.9 ± 1.2 s) (Figures 6C–6E and S6D). Replacement of with EWS1 showed the same trend, but only partly rescued the residence times (13.6 ± 1.0 s) (Figures 6B, 6D, 6E and S6B). Moreover, both the FUS and EWS-IDR fully recovered the Gal4ΔIDR search time (298 ± 21 s for Gal4ΔIDR-FUS and 275 ± 24 s for Gal4ΔIDR-EWS vs 325 ± 19 s for WT Gal4) (Figures 6F and 6G). These data indicate that IDR-mediated self-interactions enable both stable DNA binding and efficient target search. In addition, the different ability of these IDRs to restore residence and search times supports our previous observation that these forms of cooperativity are independently regulated.

**Figure 6.**
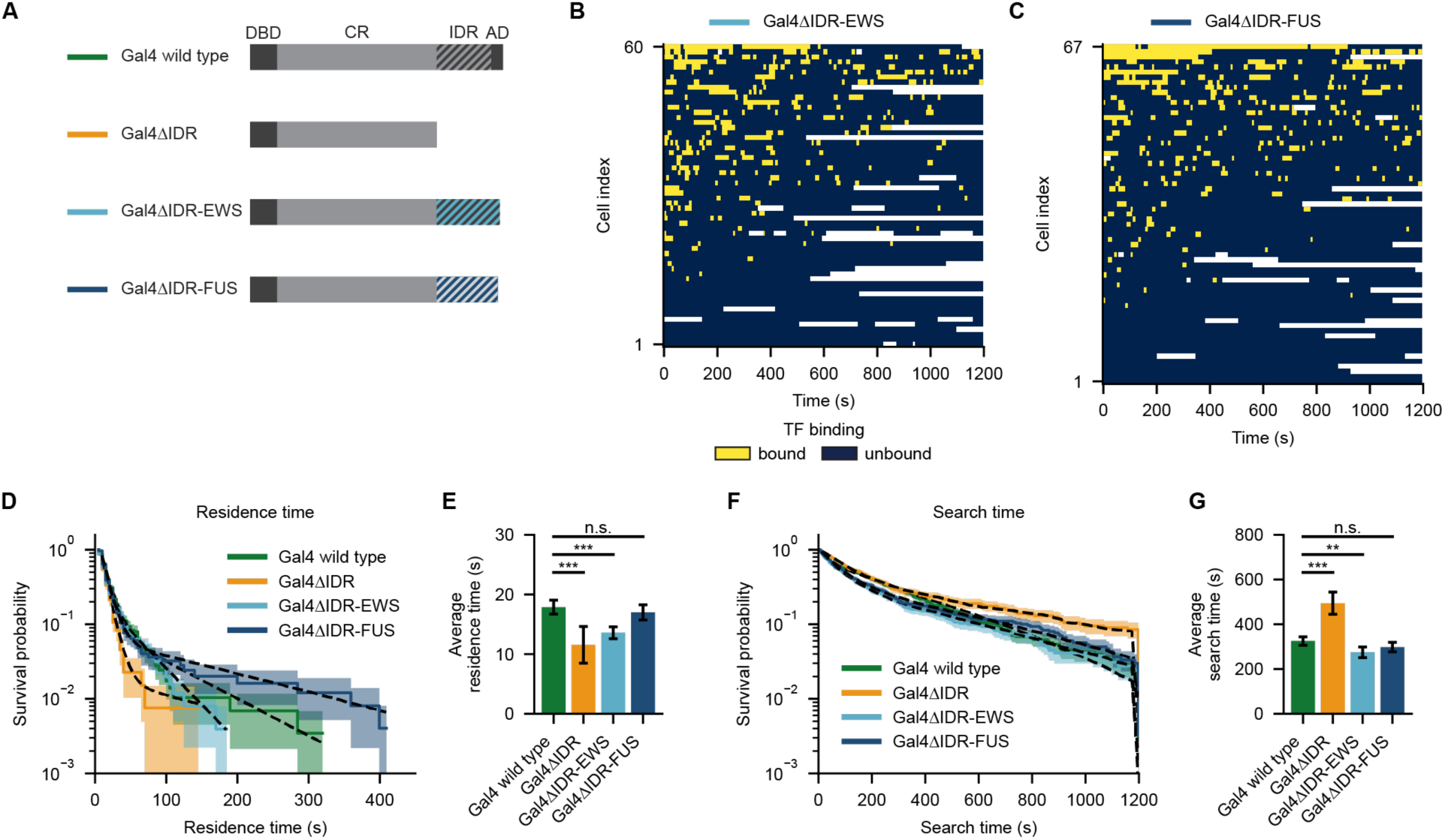
Replacement of the Gal4 IDR by mammalian IDRs rescues target binding dynamics. (A) Schematic showing 2D-representations Gal4 variants, with from top to bottom Gal4 wild type, Gal4ΔIDR, Gal4ΔIDR-EWS and Gal4ΔIDR-FUS. (B) Heatmap of binarized intensity traces capturing target search and residence time durations of single Gal4ΔIDR-EWS molecules at the *GAL* locus (blue = unbound; yellow = bound; white = no data). (C) Same as (B) for Gal4ΔIDR-FUS. (D) Survival probability distributions of residence times of Gal4 wild type (green; *n*=84 traces), Gal4ΔIDR (orange; *n*=59 traces), Gal4ΔIDR-EWS (light blue; *n*=60 traces) and Gal4ΔIDR-FUS (dark blue; *n*=67 traces) at the *GAL* locus. Solid lines indicate raw durations, shaded areas represent bootstrapped 95% confidence intervals and dotted lines show the biexponential fits. (E) Average residence times from the fits of the survival probability distributions in (D). Error bars represent 95% confidence interval from the fit. (F) Same as (D) for the search times. (G) Average search times from the fits of the survival probability distributions in (F). Error bars represent 95% confidence interval from the fit.

## DISCUSSION

In summary, we develop a unique single-molecule strategy based on our recently developed focus-feedback microscopy method,^23^ to directly capture, for the first time, both the residence and search times of single TF molecules at a specific endogenous locus in living eukaryotic cells. Our approach reveals that cooperative TF binding at neighboring TFBS support prolonged residence through multimerization and IDR self-interactions. Moreover, we discover a second form of TF cooperativity in its search process that depends on IDR-mediated self-interactions.

### Efficient TF search through IDR-mediated self-interactions

Our single-gene tracking method reveals that Gal4 search to the *GAL* genes in living cells takes ∼5 min, which is close to the diffusion limit. In search for the *GAL* locus, Gal4 encounters numerous proteins in the crowded nucleus, and needs to sample many non-specific DNA sequences before it recognizes a specific binding site. All these interactions slow down its movement compared to 3D diffusion and prolong the search time. In addition, in between two consecutive encounters at the *GAL* locus, Gal4 might associate with TFBS at other target loci, such as *GAL2*, *GAL3* and *GAL80*. Given these mechanisms that increase the search time, it is remarkable how rapid Gal4 finds the *GAL* locus, being slowed down just 4.4 times compared to the diffusion limit. The 5-min search time of Gal4 outperforms previous estimates based on conventional SMT of 5 h for the yeast TF Mbp1,^43^ 31 days for mammalian Sox2,^44^ 35 days for exogenously expressed tetR,^45^ and 465 days for the artificial Talen TF,^46^ even when compensating for the larger mammalian nucleus. Although incorrect assumptions could have biased previous estimates, comparison with our direct search time quantification shows that Gal4 target search is highly efficient.

Notably, the most commonly assumed mechanism to accelerate TF search––facilitated diffusion––is not required for Gal4 target search. In contrast to bacterial LacI, the eukaryotic TF Gal4 does not require 1D sliding and 3D hopping to find its target. Already for LacI, sliding is often non-productive, resulting in >90% sliding over its TFBS before binding, resulting in longer *in vivo* than *in vitro* search times.^7^ The blockage of TF sliding by nucleosomes may make 1D sliding in eukaryotes even less effective than in bacteria.^47^ So far, sliding of eukaryotic TFs has only been observed *in vitro*, and the gene-agnostic nature of conventional SMT has prevented testing of this mechanism *in vivo.* Our finding that 1D sliding is not required for Gal4, questions its relevance for other TFs, especially for TFs that bind to nucleosomal DNA, such as pioneering TFs.^48,49^ Nevertheless, the overall search time of Gal4 (5 min) is highly similar to bacterial LacI (3-5 min), suggesting that Gal4 must use alternative mechanisms to accelerate its search.

In this study, we find that efficient Gal4 target search requires cooperative self-interactions mediated by its C-terminal IDR. Surprisingly, we find little contribution on search from the activation domain. Loss of Gal4 activation domain does reduce transient non-specific chromatin interactions, which are generally assumed to be an intermediate step in the target search,^18,50^ but specific DNA-binding kinetics are largely unaffected. In contract, removal of the entire C-terminal IDR causes a decrease in DNA binding stability and a less efficient target search. This separation-of-function of activation domain and IDR highlights that it is not the interactions with the transcriptional machinery or the action of chromatin remodelers, but the ability of TF IDRs to self-interact that drives rapid and cooperative target recognition. The importance of cooperative self-interactions is in line with our findings that reducing the Gal4 concentration prolongs target search, while replacement of the Gal4 IDR with other self-interacting IDRs like FUS or EWS1 restores target search efficiency. Given the flexibility and dynamicity of IDRs, we propose that IDRs of DNA-bound Gal4 molecules protrude into the nucleoplasm, forming a larger target. These protruding IDRs “catch” diffusing IDR-Gal4 molecules through IDR-IDR interactions to guide them to the target locus.

For multiple yeast and mammalian TFs,^11,12,51,52^ IDRs have emerged as regions that facilitate gene-specificity together with their DBD. Mechanistically, how IDRs perform this function remains poorly understood.^53^ Our finding that IDRs directly regulate target search and binding stability provides an attractive explanation for this changed specificity, as target site selection would require compatible IDR-self interactions as well as protein-DNA interactions. Other models, where IDRs have been suggested to recognize promoter DNA,^11,15,54^ would be difficult to reconcile with our finding that exogenous FUS and EWS1 can substitute for Gal4 IDR in target search, as these IDRs would have had no evolutionary pressure to specifically recognize *GAL* promoter DNA. The IDR-swap experiments furthermore highlight that IDR-mediated self-interactions can substitute for each other and likely play a conserved role in enabling efficient target search across eukaryotic TFs.

### IDRs as regulators of multiple forms of TF cooperativity

Cooperativity is commonly observed in gene regulation by increased binding affinity at neighboring TFBS or by non-linear transcription response to TF dosage.^55^ However, without direct measurements of TF association and dissociation kinetics, most cooperativity studies and thermodynamic models do not distinguish between cooperativity in the residence time or in the search time of the transcription factor. Here, our data suggests TFs employs two mechanistically distinct forms of cooperativity that are regulated by different domains. Cooperative enhancement of TF association requires the IDR, whereas stabilization of TF residence at neighboring sites requires both the IDR and structured dimerization domains. The functions are partly separable, as both EWS and FUS restore efficient target search, yet only FUS restores prolonged residence times comparable to wild type Gal4. Notably, both forms of cooperativity are uncoupled from transcription activation, as neither IDR variant support sufficient *GAL* gene activation for growth on galactose. These observations underscore the modular yet interconnected nature of the DBD, IDR and activation domain in enabling strong transcription activation.

Cooperativity can arise through direct homotypic interactions, or indirectly via chromatin modifications, nucleosome displacement or DNA bending.^55–58^ Because removal of the activation domain has minimal impact on DNA-binding kinetics, indirect chromatin-mediated mechanisms are unlikely to explain the observed effects. DNA bending, typically mediated by the DBD,^56^ also does not account for the IDR-dependent effects. Instead, Gal4’s clustering behavior^27^ and the ability of heterologous self-interacting IDRs to restore function strongly suggest that direct self-interactions underlie both forms of cooperativity, with distinct contributions of structured and unstructured domains.

The presence of two dimerization domains suggests a potential “protein zipper” mechanism, in which sequential dimerization stabilizes DNA-bound Gal4 complexes at high-affinity TFBS. In a non-mutually exclusive manner, CR-mediated multimerization interactions between neighboring DNA-bound Gal4 dimers could promoter higher-order assemblies, further stabilized by IDR-mediated self-association. Such multimerization aligns with our previous observation that Gal4 forms clusters at target genes.^27^ Moreover, the central role of the IDR in both cooperativity forms underscore how self-interactions by the IDR are a potent mechanism to achieve cooperativity in gene regulation at multiple levels.

The prominent role of the IDR in regulating TF search and binding cooperativity suggests IDRs represent tunable modules for synthetic transcriptional activators. Although the Gal4 DBD fused to viral VP16 is widely used to activate transgenes,^33,34,59^ the DBD alone exhibits reduced stability and slower target search compared to wild-type Gal4. We predict that incorporation of the Gal4 IDR would enhance both association kinetics and binding stability in such constructs. As a proof-of-principle, fusion of various IDRs to artificial CRIPSR-VP64 activators boosted transcription activation.^60^ Understanding the cooperativity capacity by different IDRs will allow finetuning of the cooperativity as a separate regulatory layer in gene activation.

Overall, our study provides unprecedented insights into the search mechanisms of eukaryotic TFs. We invalidate the common assumption that eukaryotic TFs require facilitated diffusion for efficient target search *in vivo* and propose a mechanistic framework of how IDR-mediated self-interactions enable cooperativity in both TF association and dissociation, and thereby play a causal role in gene regulation to ensure rapid and specific transcription activation.

## METHODS

### Yeast strains and plasmids

Haploid budding yeast cells (*Saccharomyces cerevisiae*) of the BY4742 background were transformed to create the strains listed in Table S1. For all strains at least two replicates were constructed independently, which were verified by PCR and, if applicable, sequencing. All strains, plasmids and oligos used in this study are listed in Tables S1, S2 and S3, respectively. Yeast strains and plasmids are available on request.

The DNA label was introduced at either the *GAL* locus (*3′-GAL1*; YTL1852) or the *RNR2* locus (*3′-RNR2*; YTL1853), using three successive rounds of transformations, as described by Dovrat et al.^26^ Firstly, a PCR product from pTL539 encoding natMX flanked by homology arms for the 128x*tetO* array was amplified with oligo’s #1527/#1528 for integration downstream of *GAL1* or #1615/#1616 downstream of *RNR2*, followed by integration using homologous recombination and clonNAT selection. Secondly, the natMX cassette was replaced with the 128x*tetO* array using a CRISPR-Cas9-based approach^61^, where the strains were cotransformed with a URA-plasmid expressing Cas9 and a guide RNA targeting *NAT* (pTL552), together with a PCR product containing the 128x*tetO* array, amplified from plasmid pTL541 with oligo’s #1591/#1592, as double-stranded repair template, followed by removal of the Cas9 plasmid by 5-Fluoroorotic Acid (5-FOA) selection. Finally, *tetR-ymScarletI* was integrated at the *his3Δ0* locus using a single-integration vector (pTL626), digested with PacI.

To enable single-molecule imaging of Gal4 in the DNA label background, several consecutive mutations were created. First, to enable HaloTag labeling, exporter *PDR5* was deleted using the CRISPR-based approach described above, using a URA-plasmid expressing Cas9 and a guide RNA targeting *PDR5* (pTL552), together with a single-stranded oligo (#277) as repair template. Second, to create *GAL4-HaloTag-V5* in the DNA-label background (YTL1852, YTL1853), strains were transformed with a PCR product containing the *HaloTag-V5-loxP-kanMX-loxP* cassette, amplified from plasmid pTL627 with oligo’s #1202/#1203, which integrated at the 3′-end of *GAL4*.

The Gal4 binding sites at the *GAL* locus were scrambled using 2 successive rounds of transformations using the CRISPR-based approach as described before.^27^ To create the 1+1TFBS *GAL* variant (YTL1873), one of the binding sites in the *GAL7* promoter was scrambled by transforming with pTL355 and single-stranded repair oligo #832, followed by scrambling three binding sites in the *GAL1-10* promoter by transforming with pTL248 and single-stranded repair oligo #491. To create the 1TFBS (YTL1880) and the 1TFBS+*tetO* (YTL1882) *GAL* variants, both of the binding sites in the *GAL7* promoter were scrambled by transforming with pTL365 and single-stranded repair oligo #881, followed by transforming with pTL248 and either single-stranded repair oligo #491 to scramble three binding sites in the *GAL1-10* promoter in the case of the 1TFBS *GAL* variant, or single-stranded repair oligo #2126 to scramble three binding sites and add a *tetO* sequence in the case of the 1TFBS+*tetO GAL* variant.

The Gal4 promoter mutation was made using the CRISPR-based approach by transforming with pTL639 and single stranded repair oligo #1933 in the Gal4-HaloTag background (YTL1877).

Truncations and IDR-fusions of *GAL4* were created using the CRISPR-based approach described above. To create *GAL4-DBDonly* (YTL1865), pTL363 was transformed together with as repair template either the single-stranded oligo #1426 or a double-stranded PCR product, amplified from gDNA from YTL1563 with oligo’s #196/#1954. To create *GAL4ΔminiAD* (YTL1865), pTL387 was transformed together with single-stranded oligo #1648 as repair template. To create *GAL4ΔCR* (YTL1888), pTL464 was transformed together with single stranded repair oligo #2185. To create *GAL4ΔIDR* (YTL1889), *GAL4ΔIDR-EWS* (YTL1903) or *GAL4ΔIDR-FUS* (YTL1904), pTL363 was transformed together with respectively a single stranded repair oligo #2193, a double stranded repair template amplified from gBlock #2333 with #2334/#2335 or a double stranded repair template amplified from gBlock #1679 with #2331/#2332. The sequences encoding EWSR1^47–266^ and FUS^2–214^ were codon optimized for budding yeast.

For single-molecule tracking of Gal4, parent strain YTL302, containing *GAL4-HaloTag* and *pdr5::HPH*, was modified to create the truncation variants using the CRISPR-based approach as described above. Histone H3 served as control for chromatin-bound molecules, and was visualized by transforming parent strain YTL267, containing *pdr5::HPH*, with a PCR product a PCR product containing the *HaloTag-loxP-kanMX-loxP* cassette, amplified from plasmid pTL088 with oligo’s #1460/#1461, which integrated at the 3′-end of *HHT1.* In addition, the PP7-coat protein fused to GFPEnvy and a nuclear localization sequence (NLS-PCP-GFPEnvy) was integrated at the *ura3Δ0* locus by transformation with pTL174:PacI, serving as a nuclear marker during tracking experiments. The resulting strains were YTL1585 (full length Gal4-HaloTag), YTL1586 (Gal4-DBD-only-HaloTag), YTL1696 (Gal4ΔminiAD-HaloTag) and YTL1701 (HHT1-HaloTag).

### Western blot

Yeast cultures were started in synthetic complete medium containing the indicated carbon sources in the morning, diluted in the evening and grown O/N to OD_600nm_ 0.5, washed in MilliQ, pelleted and snap-frozen on dry ice. For protein extraction, cells were resuspended in 300 μL MilliQ, incubated with 300 μL 0.2M NaOH for 7 min at room temperature, centrifuged and resuspended in 500 μL 2× SDS-PAGE sample buffer (4% SDS, 20% glycerol, 0.1 M DTT, 0.125 M Tris-HCl pH 7.5 and EDTA-free protease inhibitors). Samples were incubated at 95°C for 5 min while shaking and centrifuged at 800g for 10 min at 4°C. A total of 20 μL lysate with loading buffer was run on a NuPAGE 3-8% gradient TAC gel and transferred to a 0.45-μm nitrocellulose membrane at 200 V, 1 A for 4h at 4°C. For blocking, the membrane was washed with TBS-T, incubated with PBS containing 5% milk for 1h and washed briefly with TBS-T, all at room temperature. The membrane was incubated with PBS containing 2% milk and primary antibody (1:5000) overnight at 4°C, washed three times with TBS-T for 10 min, incubated with 2% milk and secondary antibody (1:5000) for 1h at room temperature, washed three times with TBS-T for 10 min and once with PBS for 10 min, and imaged using an LI_COR Odyssey IR imager (Biosciences). Western blot analysis was performed using primary antibodies against V5 (R960-25, ThermoFisher) and tubulin (Ab6161, Abcam) and secondary antibodies Odyssey goat-anti-mouse 800 nm and Odyssey goat-anti-rat 800 nm.

The fluorescence signal of western blot images was quantified using ImageJ.^62,63^ In brief, ROIs of the same dimensions are drawn in each lane of the image. Next, a profile plot was created for each lane and a baseline was drawn manually to enclose the peak. The total area of the enclosed peak was calculated and used as a measure for the band intensity. This procedure was repeated for the signal of each primary antibody. The V5 band intensities were then normalized Tubulin bands and represented relative to the condition indicated in the figure legend.

### Growth assay

A growth assay was used to assess the galactose metabolism capacity of yeast strains, as described previously. Serial five-fold dilutions of indicated strains were spotted on YEP + 2% agarose plates, with different supplements, and growth was assessed after 3 days at 30°C. Growth on plates with 2% glucose was used as loading control. Growth on plates with 2% galactose + 20 μg/mL ethidium bromide was interpreted as functional galactose metabolism, since galactose is the only carbon source available, as the use of amino acids as carbon source is inhibited by ethidium bromide binding to mitochondrial DNA. Growth on plates with 2% raffinose + 2% galactose + 40 mM lithium chloride (LiCl) + 0.003% methionine was interpreted as non-functional galactose metabolism, as metabolism of galactose is lethal in the presence of LiCl due to the buildup of toxic metabolic intermediates. Only yeast without functional galactose metabolism can survive on these plates, using the raffinose as carbon source. The methionine is added to prevent buildup of toxic intermediates caused by LiCl inhibiting Hal2p/Met22p.^64^

### Microscopy sample preparation

Sample preparation was performed as described previously,^35,65^ with minor modifications. Yeast cells were grown in synthetic complete media supplemented with 2% raffinose and 2% galactose at 30°C. Cultures were started in the morning, diluted in the evening and grown overnight to mid-log (OD_600 nm_ 0.2–0.4). For HaloTag labeling, cultures were incubated for 15 minutes with JFX650H dye,^24^ with a concentration of 5 nM for the *GAL4* promoter mutant, 500 pM for the other Gal4-HaloTag cells & variants, and 5 pM for the H3-HaloTag cells. The cells where then washed with warm media, placed on a washed and sonicated Zeiss Coverslip HI (000000-1787-996), mounted in a Attofluor Cell Chamber (Invitrogen A7816) and immobilized using a 2% agarose pad consisting of synthetic complete media supplemented with 2% raffinose and 2% galactose, followed by focus-feedback imaging or single-molecule tracking.

### General microscopy settings

Samples were imaged on a Zeiss AxioObserver.7 / ELYRA.P1 microscope, equipped with an incubator for microscopy (Pecon Incubator XL Tirf Dark S1 Nano) set at 30°C, a Zeiss Scanning Stage Piezo 130×100 and Coherent Sapphire 405, 488, 561 and 640 nm lasers. Images were acquired with a Zeiss alpha Plan-Apochromat 100x numerical aperture (NA) 1.57 oil objective with Immersol HI 661 immersion oil and a 1.6x Zeiss optovar, yielding a pixel size of 97.09 nm. All imaging was performed using Zen Black 2.3, with highly inclined and laminated optical sheet (HILO) illumination mode and EMCCD gain set to 100x.

### Focus-feedback microscopy settings

Focus-feedback imaging was performed as described previously,^23^ with minor modifications. Images were captured with TIRF_uHP acquisition mode, a field of view of 256×256 pixels and 5 s interval for 240 time points. At every time point, the DNA label and Gal4 molecule were imaged subsequently without delay, using exposure times of 100 ms for both tracks. The tetR-ymScarlet-I DNA label (Track 1-TV2) was imaged with 561 nm excitation at ±0.0059 mW power, resulting in ±7.4 W/cm^2^ excitation intensity. The Gal4-JFX650H molecules (Track 2-TV1) were imaged with 640 nm excitation at ±0.08 mW power, resulting in ±100 W/cm^2^ excitation intensity. A filter set consisting of a Chroma ZT405/488/561rpcv2-UF1 dichroic filter and a ZET405/488/561/640mv2 emission filter was used. The emission was split in two channels (TV1 and TV2) using a Zeiss Duolink splitter holding a filter set with a Zeiss BS642 dichroic beamsplitter and Semrock BP570-650 and LP655 emission filters and imaged on two Andor EM-CCD iXon DU 897 camera’s.

To keep the DNA label in focus during the experiment (see “Focus feedback” in Methods of Pomp et al.^23^), a 300 mm cylindrical lens (Thorlabs LJ1558RM-A) was placed in a holder (Zeiss 000000-1772-065) mounted between the Duolink splitter and the front camera (imaging the DNA label), resulting in elliptical elongation of any fluorescent spot within 300 nm of the focal plane. In an identical holder between the Duolink splitter and the second camera (imaging transcription factors), a flat piece of glass as thick as the cylindrical lens was placed to keep the optical path length the same between both channels. Custom focus-feedback software, available as graphical-user interface (GUI) package (FocusFeedbackGUI), tracked the DNA label spot and used its ellipticity to adjust the *z-*position in real-time in between frames. The feedback system was calibrated using a 100 nm interval *z*-stack through 200 nm beads (TetraSpeck T7280) mounted in 1% agarose at a concentration of 2.3×10^9^ beads/ml on a Zeiss Coverslip HI coverslip. Every day prior to imaging, three *z*-stacks (101 × 100 nm) of the sample of TetraSpeck beads were acquired, which were used during the analysis to correct for the compression in the *x*-direction of images acquired by the front camera images due to the presence of the cylindrical lens and other channel misalignments (see “Channel registration” in Methods of Pomp et al.^23^).

### Focus-feedback image analysis

Localization and tracking was performed as described previously,^23^ with minor modifications. Custom python software was used to detect spots within a region of interest of 60 x 60 pixels in the center of every frame, for both the DNA label and transcription factor (TF) channels, using a Laplacian of Gaussian filter. These detections were then fitted using the L-BFGS-B fit algorithm, to determine spot intensity and localization. Next, the fitted spots were tracked over multiple frames with TrackPy,^66^ using a maximum jump distance of 6 pixels in between frames and a maximum *xy*-distance of 4 pixels (∼400 nm) between the DNA label and the TF within a frame. The choice of this 4 pixel 2D radial distance threshold was based on previously observed overlap between the same *GAL* DNA label and Gal4-EGFP clusters.^27^ We note that the *tetO* array is located downstream of *GAL1*, and thus requires a larger distance threshold compared to previously-used *PP7*-*GAL10* transcription site (threshold ∼300 nm).^23^ Frames in which the DNA label was out of focus or incorrectly tracked were excluded from the analysis.

To only include cells in which a single Gal4 molecule was labeled with JFX650H, the cell in the center of the movie was segmented using Cellpose^67^ on a sum projection of the TF channel, and in every frame the number of spot detections in the TF channel within this cell mask were counted. Cells were included only if they (i) contained at least 1 frame with 1 TF detection throughout the movie, (ii) contained no frames with more than 2 simultaneous detections, and (iii) contained at most 1 frame with 2 simultaneous detections. These selection criteria ensured exclusion of cells with multiple labeled Gal4 molecules, while preventing overfiltering of the data due to detection of noise by the spot-detection algorithm.

To discriminate between bound and unbound events, the TF intensity trace was binarized using a threshold (200 counts) (Figure 1C). The binarization threshold of 200 was based on visual spot assessments. To examine the effect of the binarization threshold on the results, the traces were binarized using a range of thresholds from 180 to 220 counts, in steps of 10. The obtained kinetic parameters were relatively stable within this threshold regime (Figure S7), showing the results are largely independent of the binarization threshold.

The binarizations were corrected by removing events lasting a single frame (1-frame gaps and jumps), as well as splitting bound events into two separate events if the relative displacement of the TF exceeded 400 nm in between frames (Figure S1B). This yielded in a clear separation between TF track intensities in bound and unbound frames, and a unimodal peak, in line with the single-molecule labeling density (Figure S1C).

Because it is unclear whether 1-frame binding events (1-frame jumps) are specifically bound to the motifs or represent non-specific binding events, transient interactions or clustered molecules, these were removed from the binarized traces. However, removal of 1-frame jumps increases search time estimates and could potentially affect comparisons between mutants if these short events occur more frequent in some conditions. To understand how this affected the analysis, the traces were binarized without removal of 1-frame jumps. Overall search times decreased but remained diffusion limited (Figures S8A and S8B), and relative differences between mutants were maintained, demonstrating that the conclusions on TF search mechanisms are not affected by removal of 1-frame jumps.

Photobleaching of Gal4 molecules could technically result in an overestimation of the search time. To estimate the degree of photobleaching, the number of bound and unbound, and total number of Gal4 molecules were plotted over time (Figures S9A and S9B). The decrease in the number of bound Gal4 molecules is likely due to selection bias during image acquisition, where cells were more often selected when a Gal4 molecule was bound to the locus (Figure S9B). The number of unbound Gal4 is stable until approximately frame 160, indicating photobleaching plays a minor role in these early time frames (Figures S9B–S9D). After frame 160, we observed a modest decrease in the number of detected Gal4 molecules, possibly from photobleaching. To check the potential effect of this photobleaching on the search time measurement, we re-calculated the search times of all experiments on only the first 160 frames of the movies, during which little photobleaching was observed. Compared to the complete time traces, truncation at 160 frames resulted in a decrease of all search times by approximately 20% (Figures S8A and S8C), but the relative differences between the mutants remained the same. We thus conclude that photobleaching affects the quantification by maximally 20%, and does not affect the presented conclusions. Because truncation of the trances resulted in loss of long events and lower statistical power, the data presented in the manuscript includes all time frames.

The distributions of bound and unbound events were plotted as survival plots, and the resulting curve was fitted by both single- and bi-exponential distributions. Remaining events lasting a single frame, not removed during binarization as they overlapped with the start/end of observation or resulted from splitting a bound event, were excluded from the fit. The survival functions are, respectively, given by:

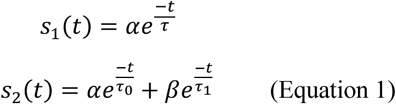

For both *s_1_*(*t*) and *s_2_*(*t*), the sum of the rate parameters *α* and *β* was normalized to 1 to extract the fractions of short and long events.

These survival functions were then corrected for the finite total time of the experiment. The observed survival function (*O_n_*, observed for *n* frames) is the result of a convolution of the real survival function with a function describing the duration of the experiment; 1 during the experiment, 0 during other times. This can be split in two parts: *A_n_* describes the part of the observed survival function due to events that started during the experiment, *B_n_* describes the part due to events that started before the experiment. With *N* the number of frames in the trace, *Δt* the time interval and *t_e_* the exposure time:

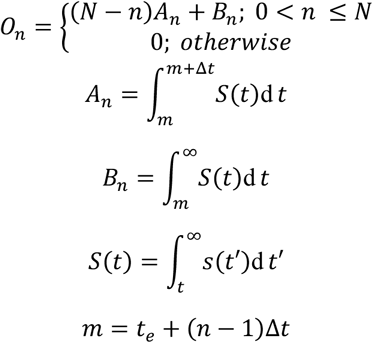

Finally normalize:

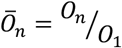

The discrete observed survival function can be derived for the distributions in Equation 1 and fitted to the data, yielding the parameters of the underlying distributions. We used the Bayesian Information Criterion (BIC) to select the best fitting model.

Even though the search time was fit best with a bi-exponential model, we noticed that the observed search times for many perturbations and for non-target *RNR2* approximated the total duration of the time traces. In addition, differences in shapes of the search time distributions complicated the comparison of the two individual exponents between different conditions. We therefore used the two-component fit to determine the average search time. The fit results of the individual exponents of the search time can be found in Table S4.

### Estimation of apparent dissociation constant

Based on *in vivo* mass spectrometry experiments, a single haploid yeast cell grown in galactose contains approximately 605 Gal4 molecules.^29^ This abundancy together with a nuclear volume of *V_nucleus_* = 2.8 × 10^−15^ L for haploid yeast grown in galactose,^68^ and Avogadro’s constant was used to calculate the nuclear Gal4 concentration:

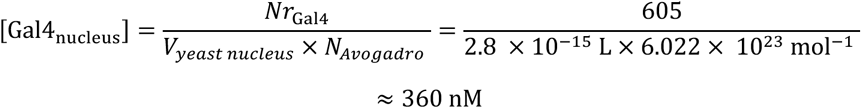

In combination with the average Gal4 residence time (*τ_on_* = 17.9 ± 1.2 s) and search time (*τ_off_* = 325 ± 19 s) obtained in this study, we estimate the apparent dissociation constant at the *GAL* locus:

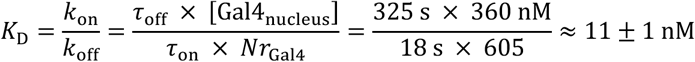

Here, we calculated the off-rate by taking the inverse of the average residence time (*τ*_on_) and corrected the on-rate (*k_on_*) for the number of available Gal4 molecules using the number (*Nr*_Gal4_) and nuclear concentration ([Gal4_nucleus_) of Gal4. The *in vivo* apparent *K*_D_ of 11 ± 1 nM represents the affinity of the *GAL* locus with 6 binding sites for full length Gal4, whereas the *in vitro* affinities of 3–35 nM were performed with one cognate TFBS and the Gal4-DBD.^28–31^ We highlight that the apparent affinities of the 1TFBS construct (*K*_D_ = 17 ± 3 nM) and Gal4-DBD-only (*K*_D_ = 41 ± 11 nM) are in the same range. Note these are only a rough estimate of the apparent *K*_D_’s, which assumes a homogenous distribution of Gal4 within the nucleus.

### Estimation of the Smoluchowski limit for diffusion-based target search

The theoretical search time, *τ*_search_, for a TF to reach its target by unhindered random 3D diffusion in a homogenous medium, can be derived using the Smoluchowski equation^32^:

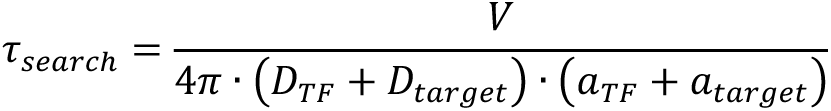

Where *V* is the volume of the medium, *D* the diffusion coefficient and *α* the size for both the TF and its target.

Using a nuclear volume of *V* = 2.8 × 10^−15^ L for haploid yeast grown in galactose,^68^ an average diffusion coefficient of *D*_target_ = 239 nm^2^/s for the locus ^23^, the measured diffusion coefficient of *D*_TF_ = 0.31 ± 0.03 µm^2^/s (Figure S2), a protein size of *a*_TF_ = 3.84 nm for Gal4 dimers,^69^ and a target site size of *a*_target_ = 0.34 nm · 17 bp, we find that the diffusion-limited search time *τ*_search_ ≈ 74 s for a single Gal4 motif. The measured average search time of 325 ± 19 s for the *GAL* locus is approximately 4.4 times longer than this theoretical limit. Although the *GAL* locus contains 6 binding sites, we chose not to correct for these 6 binding sites, because the search time for the 1TFBS construct is similar to the wild type *GAL* locus. Importantly, correcting for the number of binding sites would reduce the theoretical diffusion limit, and would make the difference between the theoretical and measured search time even larger. Similarly, even if after filtering some cells accidentally contained more than 1 labeled Gal4, correcting for this would make the difference with the theoretical diffusion limit larger. Using the effective Gal4 diffusion coefficient of the entire Gal4 population rather than of the diffusive population increases the estimated search time to ∼107 s, closer the measured search time, but still diffusion limited. In all these cases, the conclusion that Gal4 search is slower than the Smoluchowski limit thus remains valid. We note that the Smoluchowski limit describes the first possible Gal4-target encounter by 3D diffusion, but does not consider the orientation of the molecules.

### Single-molecule tracking microscopy settings

Single-molecule tracking (SMT) was performed as described previously,^35^ with minor modifications. Imaging was performed using TIRF_uHP acquisition mode. Conventional SMT data was acquired using a field of view of 256×256 pixels, 30 ms exposure time and 200 ms interval. Fast SMT data was acquired using a field of view of 128×128 pixels, 12 ms exposure time and continuous acquisition. At every time point, the Gal4 molecules and nuclear PP7 coat protein (PCP) were imaged simultaneously. The Gal4-JFX650H molecules (TV1) were imaged with 640 nm excitation at ±0.60 mW power, resulting in ±750 W/cm^2^ excitation intensity. The nuclear PCP-GFPenvy (TV2) was imaged with 488 nm excitation at ±0.0069 mW power, resulting in ±8.6 W/cm^2^ excitation intensity. A Zeiss filter set consisting of an MBS 405/488/642 dichroic filter and a LBF 405/488/642 emission filter was used. The emission was split in two channels (TV1 and TV2) using a Zeiss duolink splitter holding a filter set with a Zeiss BS642 dichroic beamsplitter and Semrock BP495–550 and LP655 emission filters and imaged on two Andor EM-CCD iXon DU 897 cameras.

### Single-molecule tracking image analysis

Analysis of single-molecule tracking data was performed as described previously,^35^ with minor modifications. Single-molecule tracking movies were analyzed using a custom MATLAB software based on MatlabTrack v6.0 package. For each movie, a max projection of the PCP-GFPenvy signal was used to segment the nuclei, creating regions of interest for the analysis in which Gal4 molecules were tracked. Bound molecules were selected based on a minimal track length of 4 frames, a maximum frame-to-frame jump of 0.22 μm and a maximum end-to-end distance of 0.35 μm. The cumulative distribution of residence times of bound Gal4 molecules was corrected for photobleaching based on the photobleaching kinetics of the bound histone H3 population.^20^ The resulting photobleaching-corrected survival distributions were fitted with a double-exponential distribution to obtain the fractions of short- and long-bound molecules.

The diffusion coefficients of Gal4-HaloTag (YTL1585) and H3-HaloTag (YTL1701) were estimated from the fast SMT data. For every track, the mean square displacement (MSD) of the smallest N time lags was determined from which the diffusion coefficient was determined using the equation below, according to Michalet^70^:

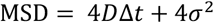

Here, *D* is the diffusion coefficient, *Δt* the time delay between frames, and *σ* the precision of the localization uncertainty. The diffusion coefficient distributions of individual replicates of Gal4 and H3 were fit with bimodal and unimodal distributions, respectively. The fast component of Gal4, representing diffusing molecules, was used for the calculation of the Smoluchowski-limit.

### Structure prediction

For the structure prediction using Alphafold3,^39^ two full length Gal4 peptide sequences, 4 Zn^2+^ ions and the sense & antisense sequence of TFBS 3 in the *GAL1-10* promoter (‘5-CGGAAGACTCTCCTCCG-3’ and ‘5-CGGAGGAGAGTCTTCCG-3’) were used as input. The zip file containing the Alphafold3 result was used as input for AlphaBridge high confident interaction interfaces.^40^

### Quantification and statistics

Focus-feedback imaging data was taken at least three biological replicates over multiple days. The total number of traces per condition are shown in the heatmaps. *P* values for comparisons between fit parameters displayed in bar graphs were determined using Mann Whitney *U test* and significance is defined as: not significant (n.s), *P* > 0.05; \**P* < 0.05, \*\**P* < 0.01, \*\*\**P* < 0.001.

Error bars of fit parameters show 95% confidence intervals. Western blot quantification displayed in bar graphs show fold change compared to wild type from at least two biological replicates with error bars indicating the standard deviation.

## ACKNOWLEDGMENTS

We thank A. Coulon, Curie Institute, A.S. Hansen, MIT, R.G.H. Lindeboom, NKI, B. van Steensel, NKI, T.R. Brummelkamp, NKI, N. Scholes, NKI, A.R. Krebs, EMBL, and members of the TLL lab for helpful feedback and suggestions. We thank Luke Lavis for the dye JFX650. We thank the Research High Performance Computing Facility of the NKI for assistance. This work was supported by an institutional grant of the Dutch Cancer Society and of the Dutch Ministry of Health, Welfare and Sport, Oncode Institute, which is partly financed by the Dutch Cancer Society, the Dutch Research Council (VIDI: VI.Vidi.213.031) and the European Research Council (ERC Starting Grant 755695 BURSTREG).

## AUTHOR CONTRIBUTIONS

Conceptualization: JVWM, TLL; Methodology: JVWM, WP, DM; Investigation: JVWM, WJJ, WP; Funding acquisition: TLL; Supervision: TLL; Writing – original draft: JVWM, TLL; Writing – review & editing: JVWM, WP, DM, WJJ, TLL.

## SUPPLEMENTAL INFORMATION

**Figure S1.**
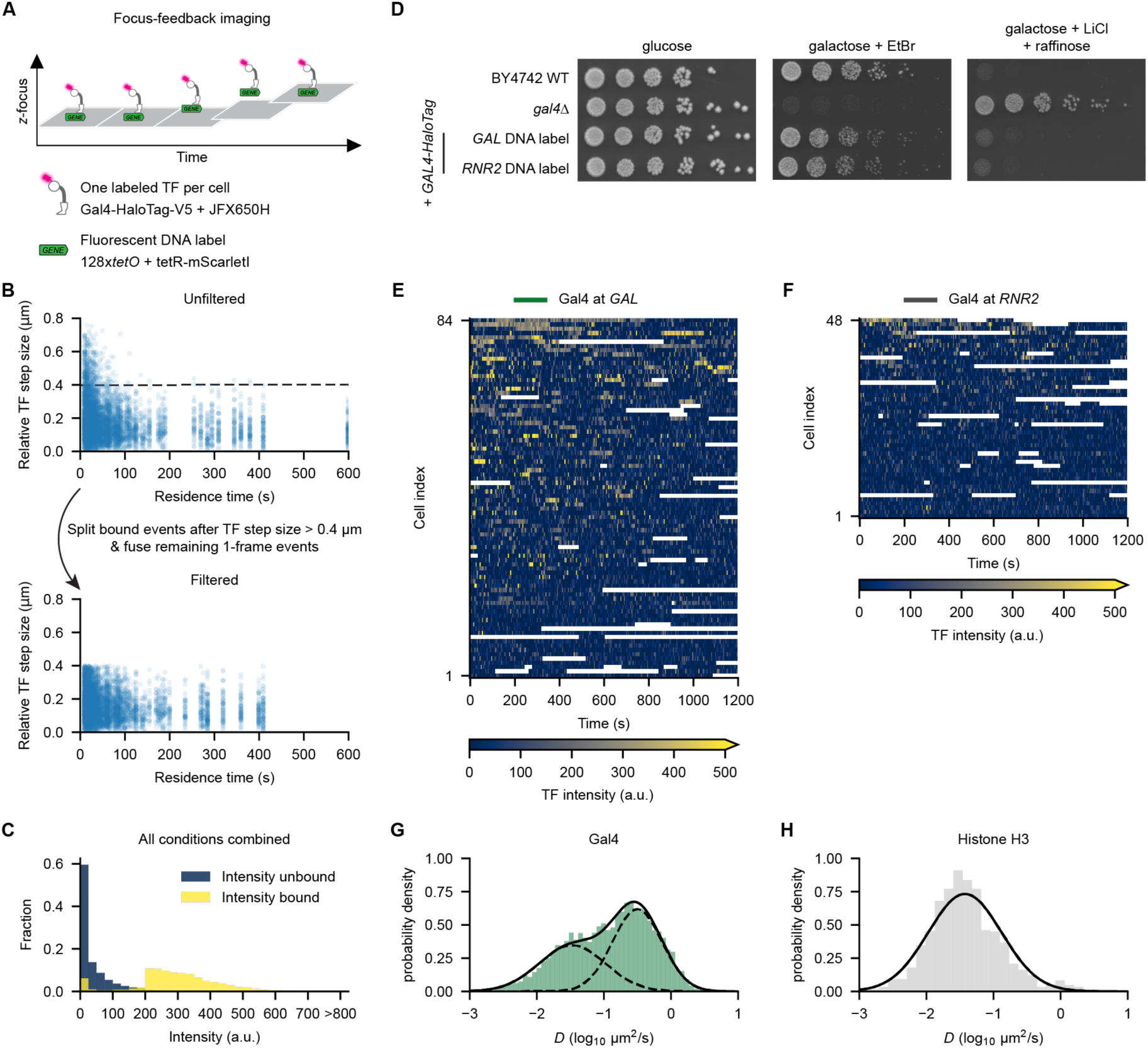
Workflow of single-molecule focus feedback imaging to quantify Gal4 DNA-binding dynamics at the *GAL* target locus. (A) Schematic of focus feedback microscopy method which enables prolonged imaging of a single locus by refocusing on the proximal DNA label, while capturing the binding kinetics of a single labeled TF molecule^23^. (B) Scatterplots of relative TF step sizes between frames, sorted by the residence time of the binding event they belonged to, from all conditions combined. Shown are TF step size versus residence time after initial binarization (top) and after splitting bound events at steps > 0.4 μm and correcting for 1-frame events (bottom). (C) Histogram of intensities from all unbound events (blue) and bound events (yellow), from all conditions combined. (D) Growth assay of indicated strains to assess the effect of genomic alterations at *GAL* and *RNR2* on their galactose metabolism capability. Shown are 5-fold serial dilutions on YEP + 2% glucose (dilution control), YEP + 2% galactose + 20 μg/mL ethidium bromide (growth = functional galactose metabolism) and YEP + 2% raffinose + 2% galactose + 40 mM lithium chloride + 0.003% methionine (no growth = functional galactose metabolism). As expected, insertion of the DNA-label and fusion of the HaloTag-V5 to Gal4 did not affect growth on galactose. (E) Heatmap of raw intensity traces capturing binding of single wild type Gal4 molecules at the *GAL* target locus. White indicates no data. (F) Same as (E) but at the *RNR2* locus. (G, H) Probability density histograms for the diffusion coefficient from combining 5 replicates of (G) wild type Gal4 (*n*=4442 traces from 195 movies) and (H) 2 replicates of histone H3 (grey, *n=*1644 traces from 79 movies), estimated from continuous and fast single-molecule tracking with 12 ms interval. The data of individual replicates were fit with a bimodal distribution (solid line, unimodal fits in dotted lines) for Gal4 and a unimodal distribution for histone H3. The slow component of Gal4 (*D* = 0.034 ± 0.005 μm^2^/s) shows similar diffusion as histone H3 (*D* = 0.038± 0.003 μm^2^/s), indicating chromatin bound Gal4 molecules. The fast component of Gal4 represents diffusing Gal4 (*D* = 0.31 ± 0.03 μm^2^/s) and was used for the calculation of the Smoluchowski-limit. Errors indicate standard deviations of the replicates.

**Figure S2.**
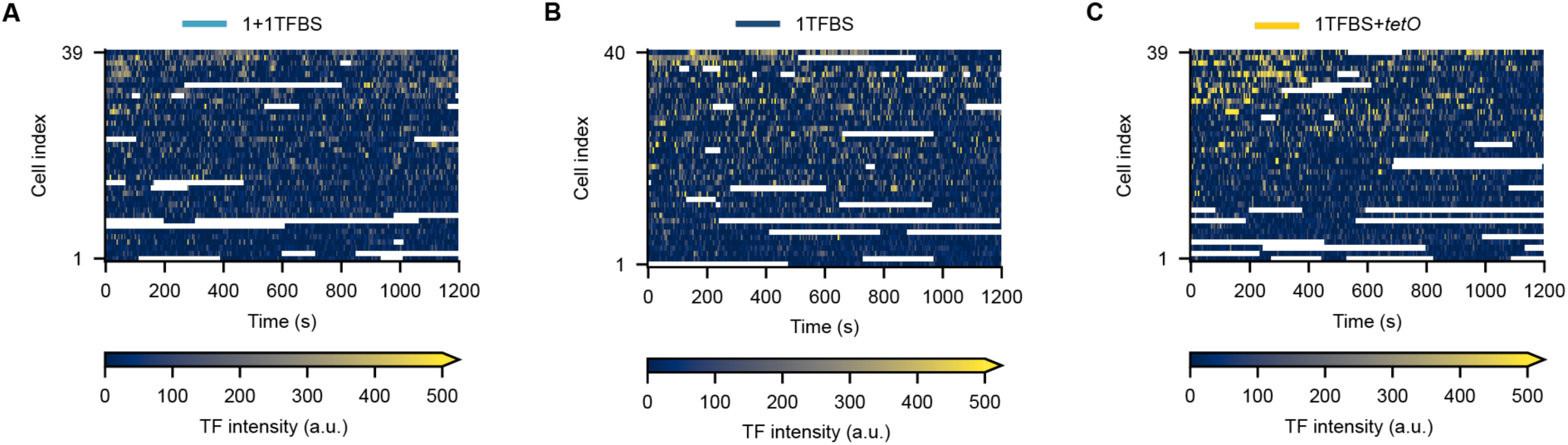
Traces of Gal4 binding at *GAL* locus variants with less binding sites. (A) Heatmap of raw intensity traces capturing target binding of single Gal4 molecules at the 1+1TFBS variant of the *GAL* locus. White indicates no data. (B, C) Same as (A) for the 1TFBS (B) or 1TFBS+*tetO* (C) variants of the *GAL* locus.

**Figure S3.**
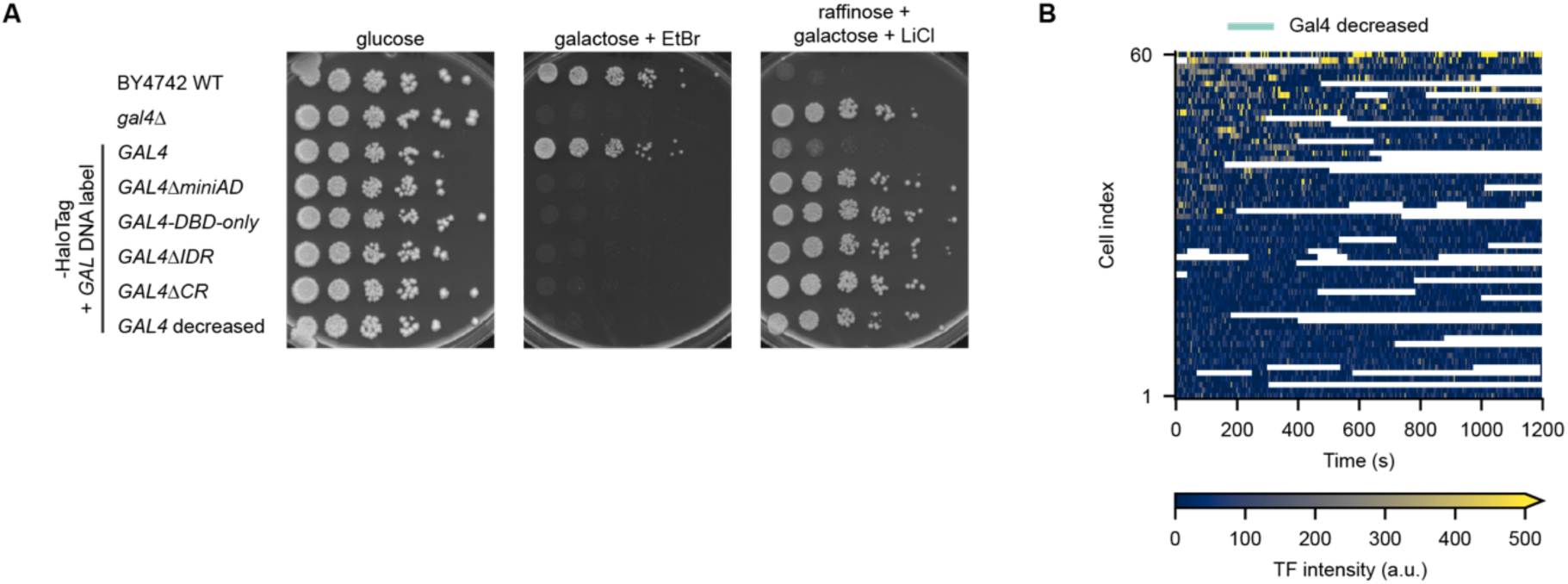
Characterization of Gal4 mutant strains used in this study. (A) Growth assay of indicated strains to assess the effect of Gal4 perturbations on their galactose metabolism capability. Shown are 5-fold serial dilutions on YEP + 2% glucose (dilution control), YEP + 2% galactose + 20 μg/mL ethidium bromide (growth = functional galactose metabolism) and YEP + 2% raffinose + 2% galactose + 40 mM lithium chloride + 0.003% methionine (no growth = functional galactose metabolism). All Gal4 mutants impede growth on galactose, comparable to the *gal4*Δ strain. (B) Heatmap of raw intensity traces capturing target search of single Gal4 molecules at the *GAL* locus in a background with decreased Gal4 protein levels. White indicates no data.

**Figure S4.**
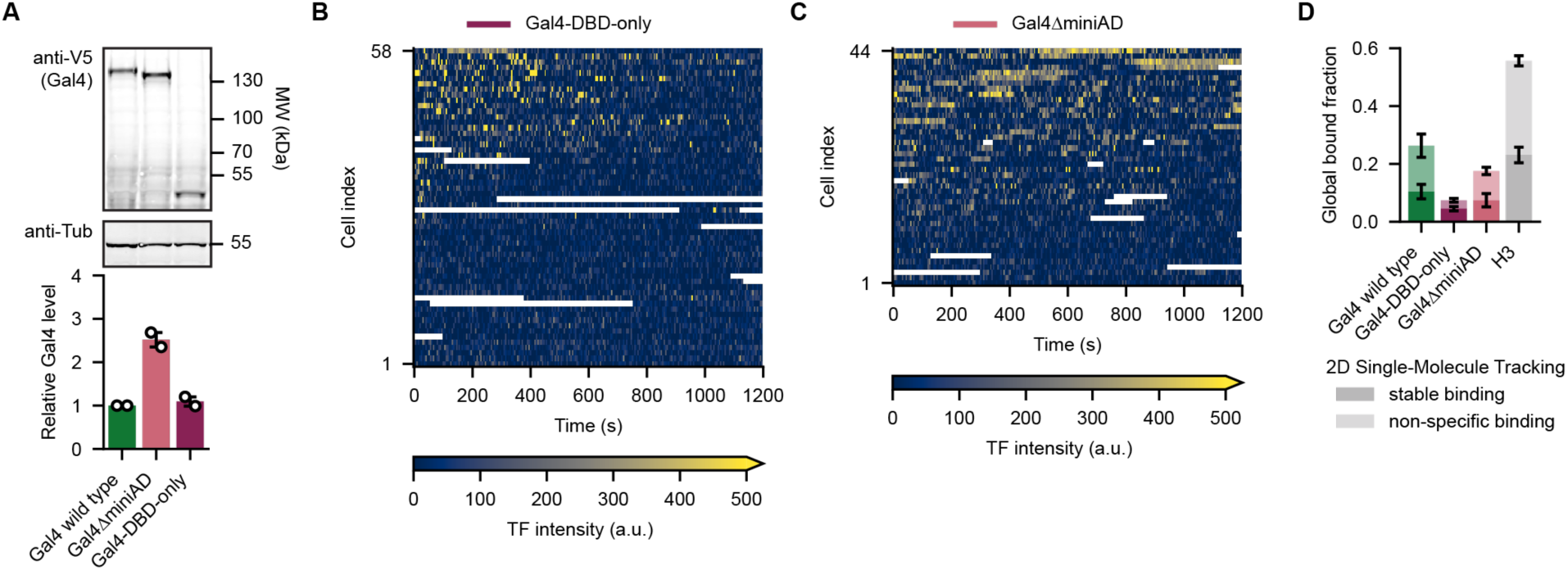
Characterization of Gal4-DBD-only and Gal4ΔminiAD mutant strains. (A) Assessment of protein levels by western blot (top) and quantification (bottom) of Gal4 wild type (green), Gal4ΔminiAD (pink) and Gal4-DBD-only (dark pink) mutants in the HaloTag-V5 background. Western blot image is a representative example of two independent experiments. The anti-V5 antibody signals are normalized to anti-Tubulin and to the expression of Gal4 wild type, resulting in the bar graph, in which open circles represent the results of individual replicate experiments (*n=*2), colored bars indicate their mean and error bars the standard deviation. (B) Heatmap of raw intensity traces capturing target search of single Gal4-DBD-only molecules at the *GAL* locus. White indicates no data. (C) Same as (B) for traces of Gal4ΔminiAD. (D) Conventional single-molecule tracking with 200 ms interval, showing the fraction of specific (stable multiple second binding, dark) and transient non-specific binding (∼1 s, light), as determined by fitting the residence time distribution of bound molecules, for indicated mutants and histone H3 as a control. Error bars indicate SEM of 3 replicates for Gal4 wild type (*n=*1048 traces), 7 for Gal4-DBD-only (*n=*957 traces), 3 for Gal4ΔminiAD (*n=*1956 traces) and 4 for histone H3 (*n=*1999 traces).

**Figure S5.**
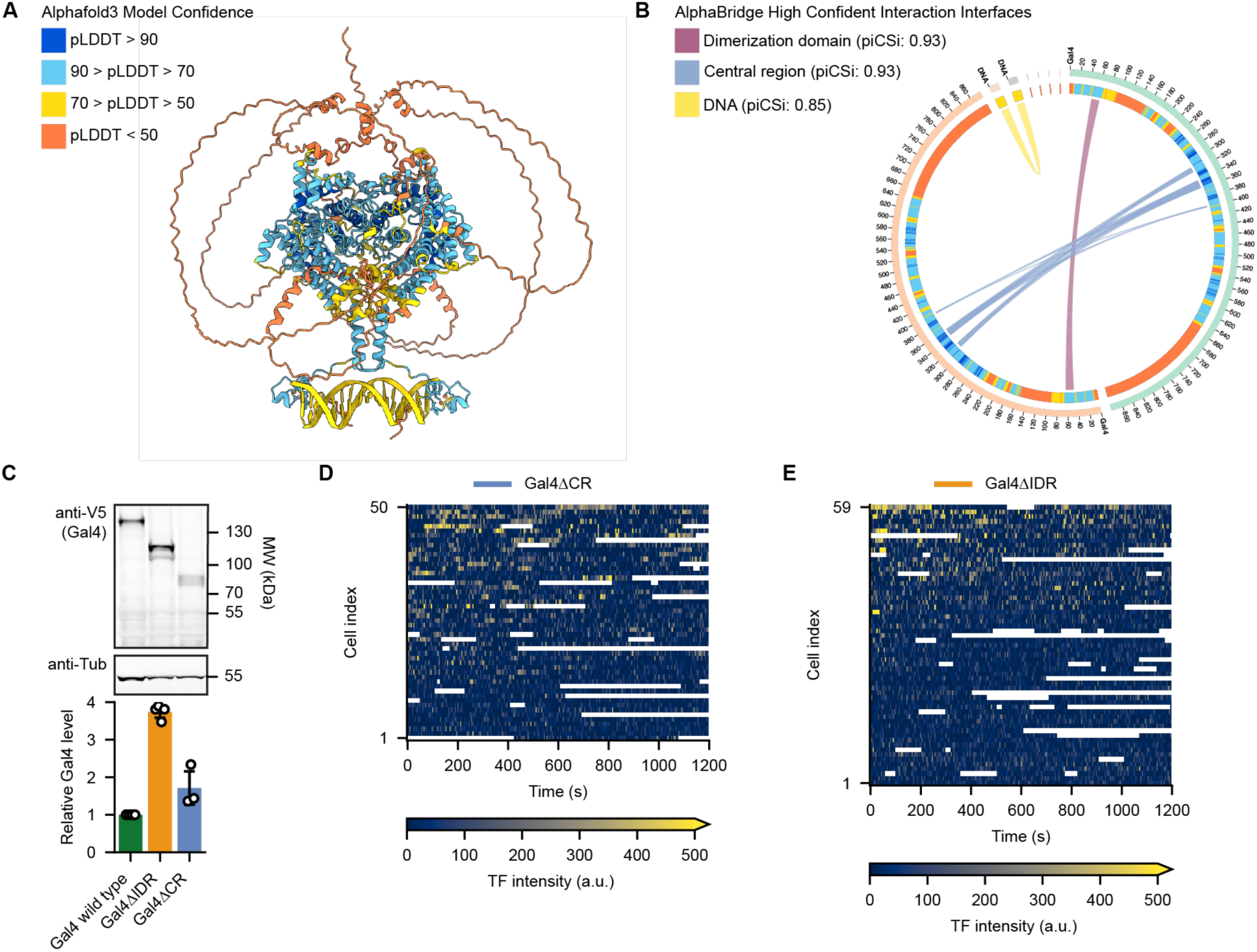
Prediction of Gal4 structure and characterization of Gal4ΔCR and Gal4ΔIDR mutant strains. (A) Alphafold3 model of two full-length Gal4 molecules with DNA (sense + antisense sequence of TFBS 3 in the *GAL1-10* promoter) 4 Zn^2+^ ions.^39^ The color of the structure shows the pLDDT scores, indicating the confidence of the predicted model. (B) Predicted high confident interaction interfaces from the Alphafold3 model by AlphaBridge.^40^ Interaction piCSi confidence scores are indicated. As expected, the two DNA strands (yellow) and the Gal4 dimerization domains (dark pink) interact. Interestingly, a strong interface is predicted between the structured domains in the Gal4 central region (purple). (C) Assessment of protein levels by western blot (top) and quantification (bottom) of Gal4 wild type (green), Gal4ΔIDR (orange) and Gal4ΔCR (purple) mutants in the HaloTag-V5 background. Western blot image is a representative example. The anti-V5 antibody signals are normalized to anti-Tubulin and to the expression of Gal4 wild type, resulting in the bar graph, in which open circles represent the results of individual replicate experiments (*n*=4 for wild type and Gal4ΔIDR, *n*=3 for Gal4ΔCR), colored bars indicate their mean and error bars the standard deviation. (D) Heatmap of raw intensity traces capturing target search of single Gal4ΔCR molecules at the *GAL* locus. White indicates no data. (E) Same as (D) for traces of Gal4ΔIDR.

**Figure S6.**
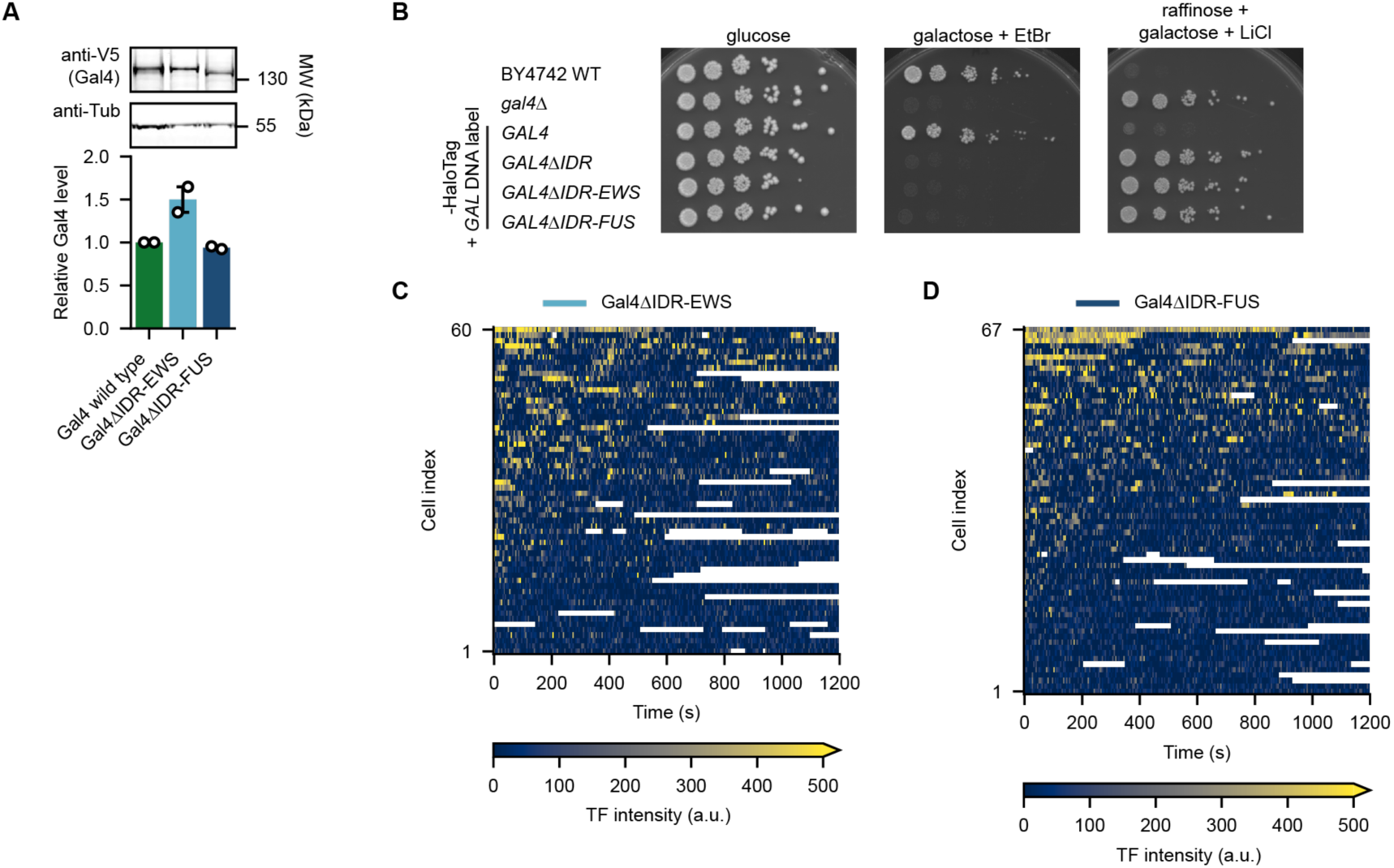
Characterization and imaging of Gal4ΔIDR-EWS and Gal4ΔIDR-FUS mutant strains. (A) Assessment of protein levels by western blot (top) and quantification (bottom) of Gal4 wild type (green), Gal4ΔIDR-EWS (light blue) and Gal4ΔIDR-FUS (dark blue) mutants in the HaloTag-V5 background. Western blot image is a representative example of two independent experiments. The anti-V5 antibody signals are normalized to anti-Tubulin and to the expression of Gal4 wild type, resulting in the bar graph, in which open circles represent the results of individual replicate experiments (*n*=2), colored bars indicate their mean and error bars the standard deviation. (B) Growth assay of indicated strains to assess the effect of Gal4 perturbations on their galactose metabolism capability. Shown are 5-fold serial dilutions on YEP + 2% glucose (dilution control), YEP + 2% galactose + 20 μg/mL ethidium bromide (growth = functional galactose metabolism) and YEP + 2% raffinose + 2% galactose + 40 mM lithium chloride + 0.003% methionine (no growth = functional galactose metabolism). All Gal4 mutants impede growth on galactose, comparable to the *gal4*Δ strain. (C) Heatmap of raw intensity traces capturing target search of single Gal4ΔIDR-EWS molecules at the *GAL* locus. White indicates no data. (D) Same as (C) for traces of Gal4ΔIDR-FUS.

**Figure S7.**
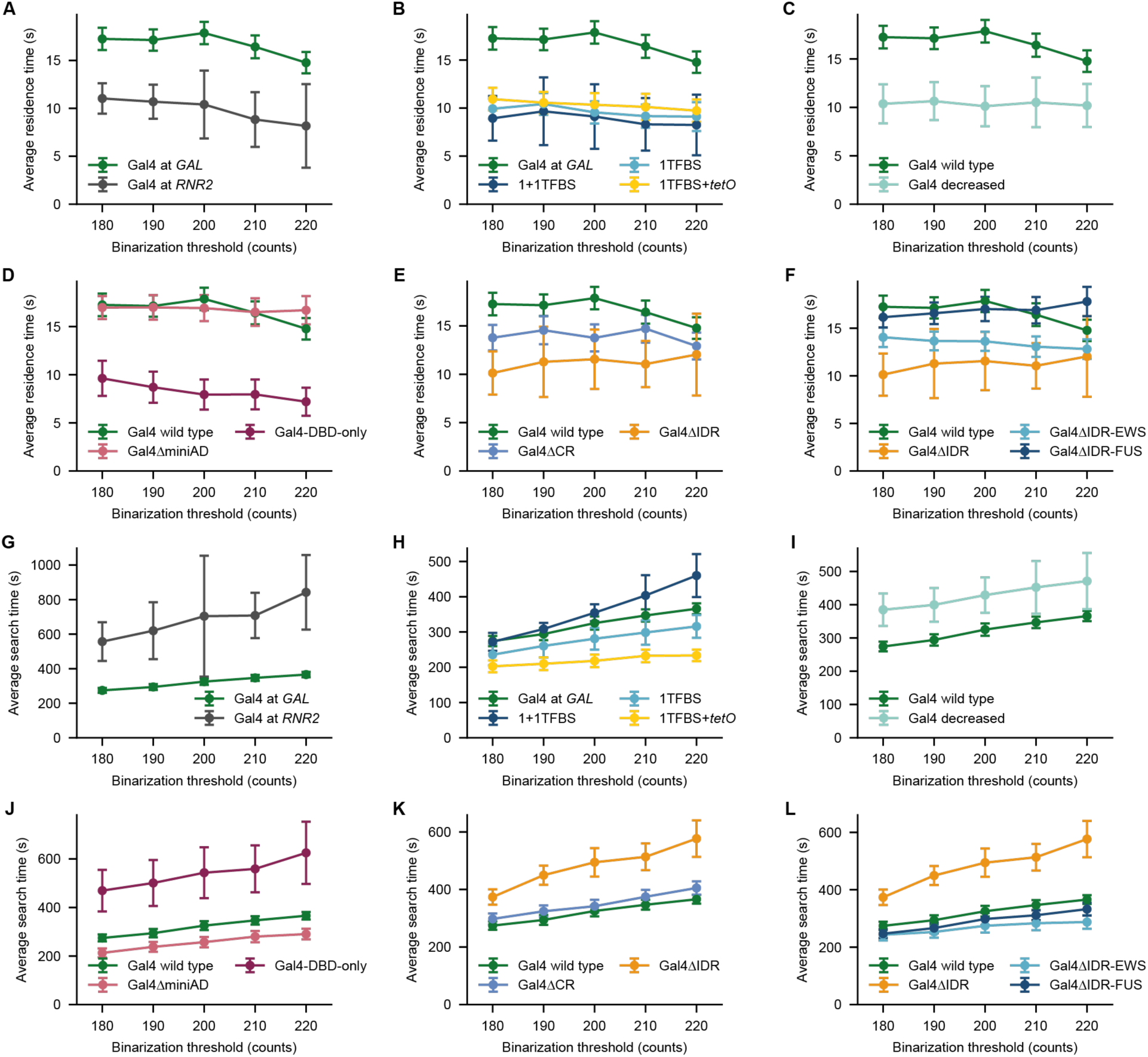
TF kinetics are largely independent of binarization threshold. (A–F) Average residence times calculated using a range of binarization thresholds (180–220 counts, with steps of 10), demonstrating that relative differences between conditions remain valid across a range of thresholds. Panels (A–F) correspond to Figures 1-6, respectively. Error bars represent 95% confidence interval from the fit. (G–L) Same as (A–F) for average search times. Panels (G–L) correspond to Figures 1–6, respectively.

**Figure S8.**
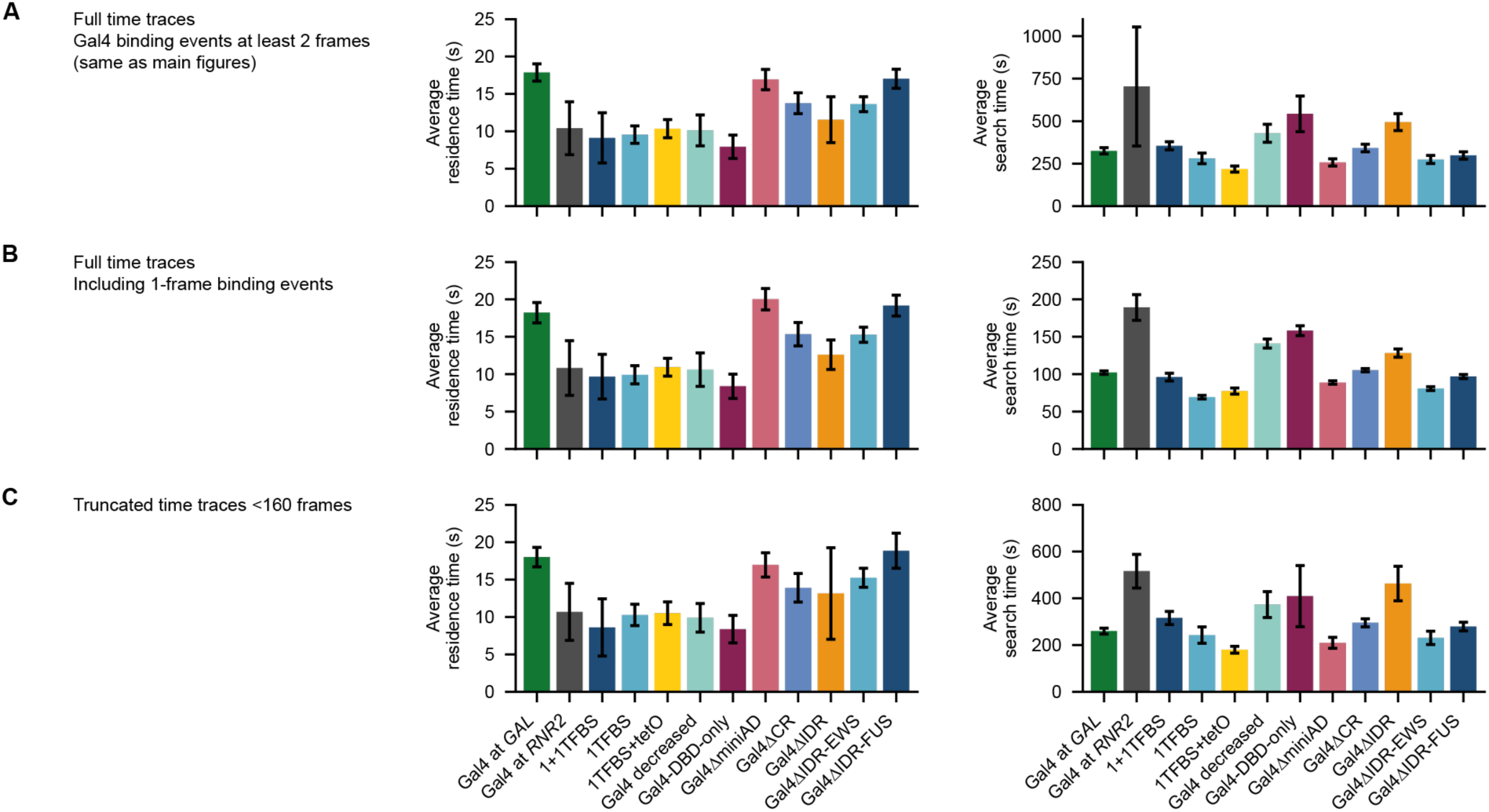
Relative TF kinetics are unaffected by removal of 1-frame binding events or photobleaching. (A) Average residence and search time parameters from the fits (mean ± 95% CI) from processed binarized traces as presented in the main figures. (B) Same as (A) except that the binarized traces are processed without the removal of 1-frame binding events. Although the average search times decrease because of these 1-frame TF-DNA label colocalizations, they remain diffusion limited. (C) Same as (A) except that only the first 160 frames of the traces are considered, which are unaffected by photobleaching. Although the average search times decrease, relative differences between the mutants remained the same.

**Figure S9.**
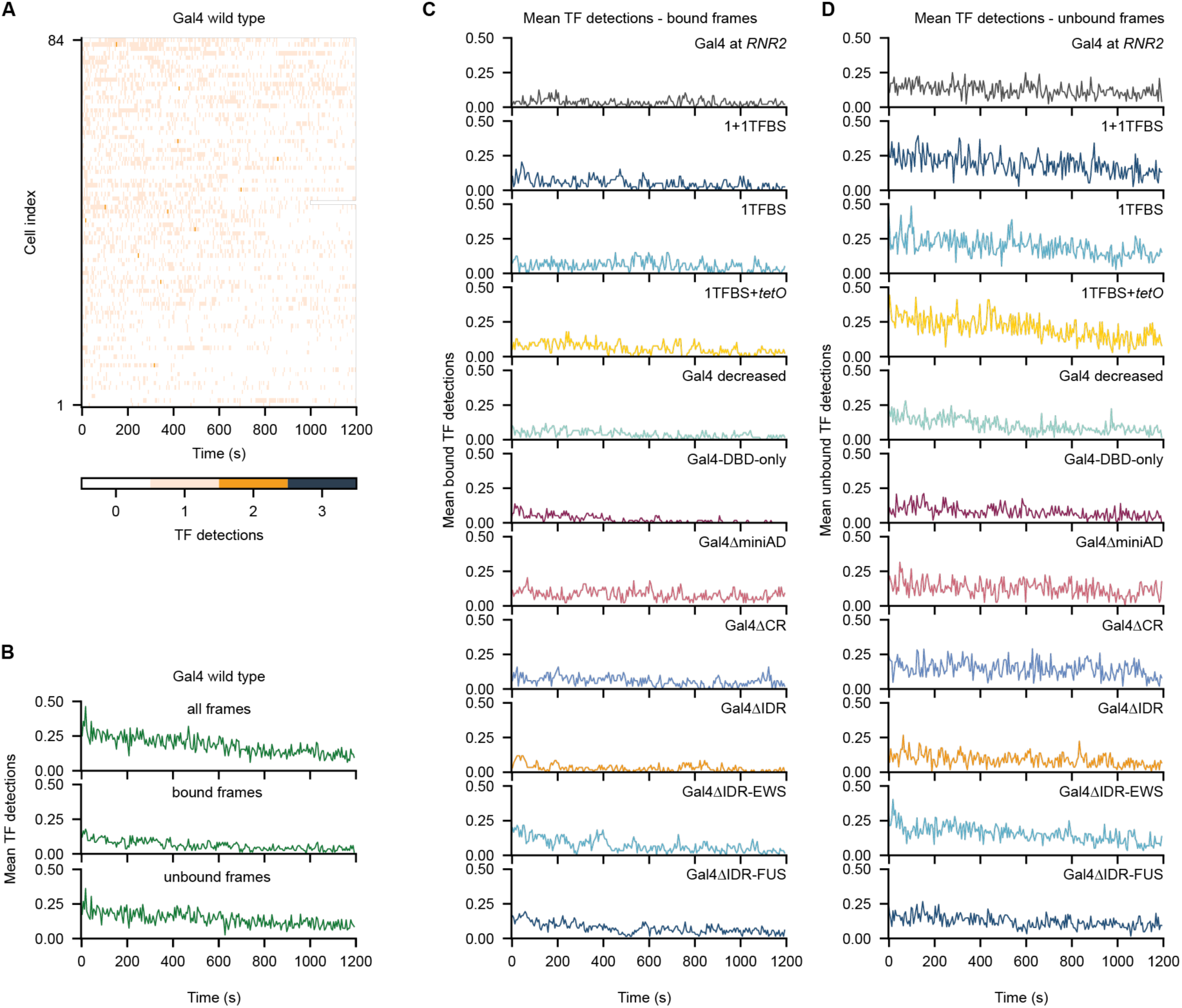
Single-molecule focus-feedback imaging leads to minimal photobleaching. (A) Heatmap of number of TF spot detections within the tracked cell over time for focus-feedback imaging of wild type Gal4 molecules at the *GAL* target locus. (B) Average TF spot detections over time from heatmap in (A), including spot detections in all frames (top), spot detections in bound states normalized by bound+unbound frames (middle), and spot detections in unbound states normalized by unbound frames only (bottom). (C) TF spot detections in bound states, normalized by bound+unbound frames, for all other experimental conditions. (D) TF spot detections in unbound states, normalized by unbound frames only, for all other experimental conditions.

**Video S1**

Example movie of a yeast cell with *GAL* DNA label (left; green) which is tracked in 3D using active focus feedback and simultaneous detection of binding of a single Gal4 molecule (center; magenta), together with the merge (right). Squares indicate the center of the tracked spots for both the DNA label and TF channel. The total duration of the movie is 20 minutes, with 5 seconds between every frame.

**Table S1.** Yeast strains used in this study.

| Strain | Genotype | Source |
| --- | --- | --- |
| BY4742 | MATα his3Δ1 leu2Δ0 lys2Δ0 ura3Δ0 | Euroscarf |
| YTL559 | BY4742 with *gal4Δ* | 27 |
| DNA label + HaloTag strains |  |  |
| YTL1852 | BY4742 with tetOx128 3'-GAL + his3Δ1:tetR1-ymScarletI (pTL626) + pdr5Δ + GAL4-HaloTag-V5 | This study |
| YTL1853 | BY4742 with tetOx128 3'-RNR2 + his3Δ1:tetR1-ymScarletI (pTL626) + pdr5Δ + GAL4-HaloTag-V5 | This study |
| YTL1865 | BY4742 with tetOx128 3'-GAL + his3Δ1:tetR1-ymScarletI (pTL626) + pdr5Δ + GAL4(Δ95-881)-HaloTag-V5 | This study |
| YTL1869 | BY4742 with tetOx128 3'-GAL + his3Δ1:tetR1-ymScarletI (pTL626) + pdr5Δ + GAL4(Δ840-881)-HaloTag-V5 | This study |
| YTL1873 | BY4742 with tetOx128 3'-GAL + his3Δ1:tetR1-ymScarletI (pTL626) + pdr5Δ + GAL4-HaloTag-V5 + 1x scrUAS 5'GAL7 + 3x scrUAS 5'GAL10 | This study |
| YTL1877 | BY4742 with tetOx128 3'-GAL + his3Δ1:tetR1-ymScarletI (pTL626) + pdr5Δ + pGAL4mut-GAL4-HaloTag-V5 | This study |
| YTL1880 | BY4742 with tetOx128 3'-GAL + his3Δ1:tetR1-ymScarletI (pTL626) + pdr5Δ + GAL4-HaloTag-V5 + 2x scrUAS 5'GAL7 + 3x scrUAS 5'GAL10 | This study |
| YTL1882 | BY4742 with tetOx128 3'-GAL + his3Δ1:tetR1-ymScarletI (pTL626) + pdr5Δ + GAL4-HaloTag-V5 + 2x scrUAS 5'GAL7 + 1x tetO & 3x scrUAS 5'GAL10 | This study |
| YTL1888 | BY4742 with tetOx128 3'-GAL + his3Δ1:tetR1-ymScarletI (pTL626) + pdr5Δ + GAL4(Δ95-650)-HaloTag-V5 | This study |
| YTL1889 | BY4742 with tetOx128 3'-GAL + his3Δ1:tetR1-ymScarletI (pTL626) + pdr5Δ + GAL4(Δ651-881)-HaloTag-V5 | This study |
| YTL1903 | BY4742 with tetOx128 3'-GAL + his3Δ1:tetR1-ymScarletI (pTL626) + pdr5Δ + GAL4(Δ95-881)-HaloTag-V5 | This study |
| YTL1904 | BY4742 with tetOx128 3'-GAL + his3Δ1:tetR1-ymScarletI (pTL626) + pdr5Δ + GAL4(Δ95-881)-HaloTag-V5 | This study |
| HaloTag strains |  |  |
| YTL267 | BY4742 with *pdr5::HPH* | 71 |
| YTL1563 | BY4742 with pdr5::HPH + GAL4(Δ95-881)-HaloTag w/o kanMX | This study |
| YTL1585 | BY4742 with pdr5::HPH + GAL4-HaloTag w/o kanMX + ura3Δ0:PCP-GFPenvy (pTL174) | This study |
| YTL1586 | BY4742 with pdr5::HPH + GAL4(Δ95-881)-HaloTag w/o kanMX + ura3Δ0:PCP-GFPenvy (pTL174) | This study |
| YTL1696 | BY4742 with pdr5::HPH + GAL4(Δ840-881)-HaloTag w/o kanMX + ura3Δ0:PCP-GFPenvy (pTL174) | This study |
| YTL1701 | BY4742 with pdr5::HPH + HHT1-HaloTag w/ kanMX + ura3Δ0:PCP-GFPenvy (pTL174) | This study |

**Table S2.**
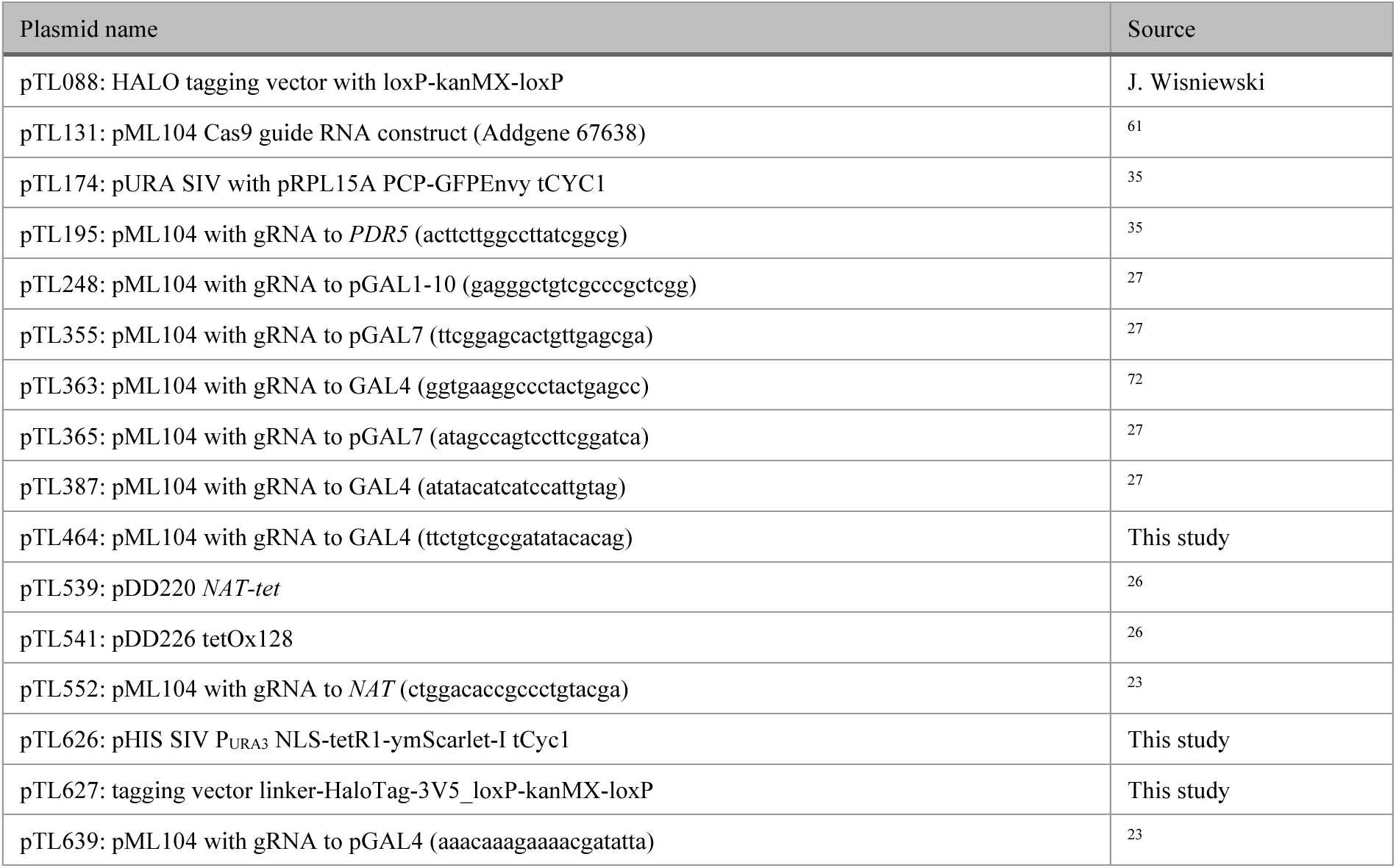
Plasmids used in this study.

**Table S3.**
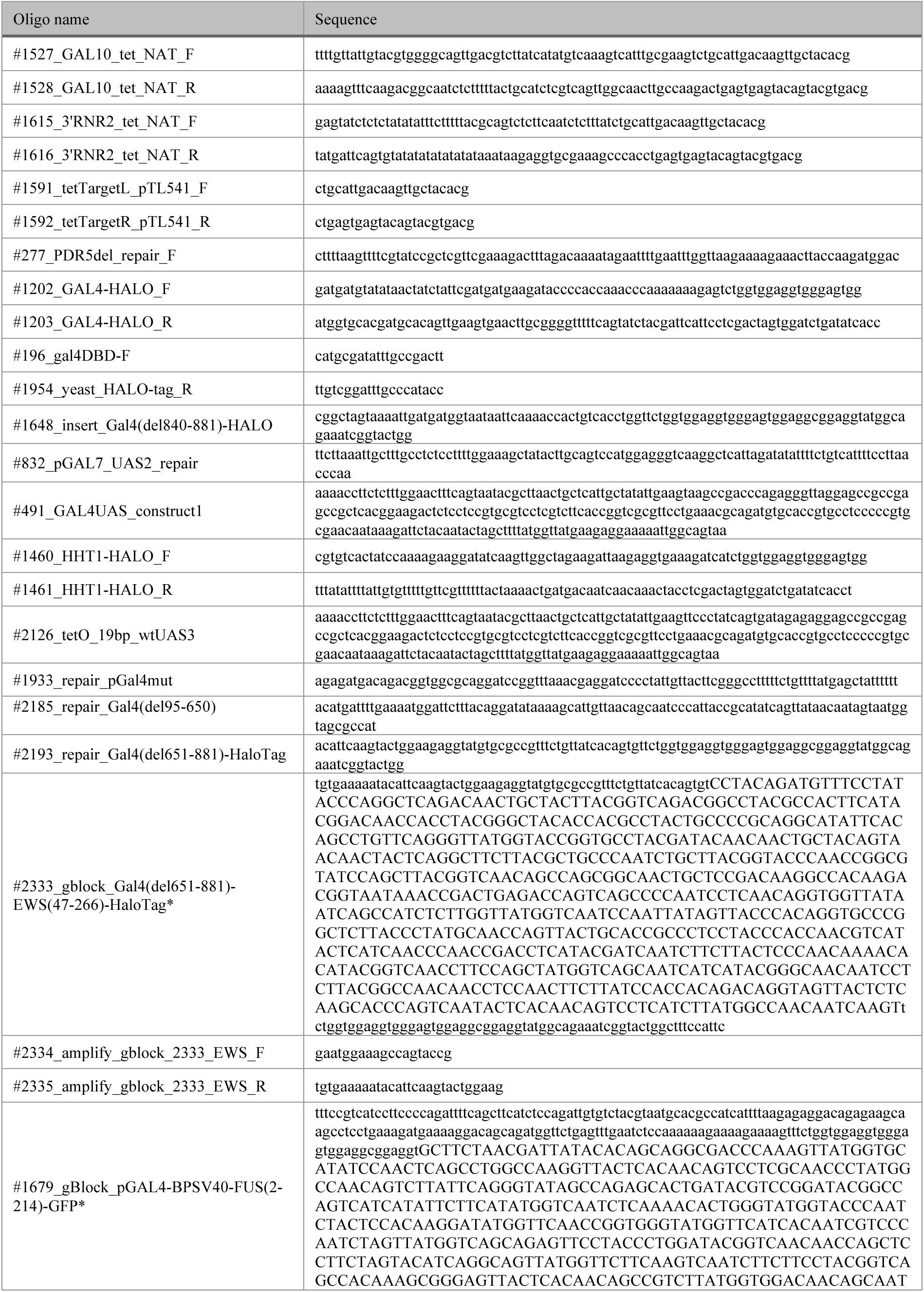
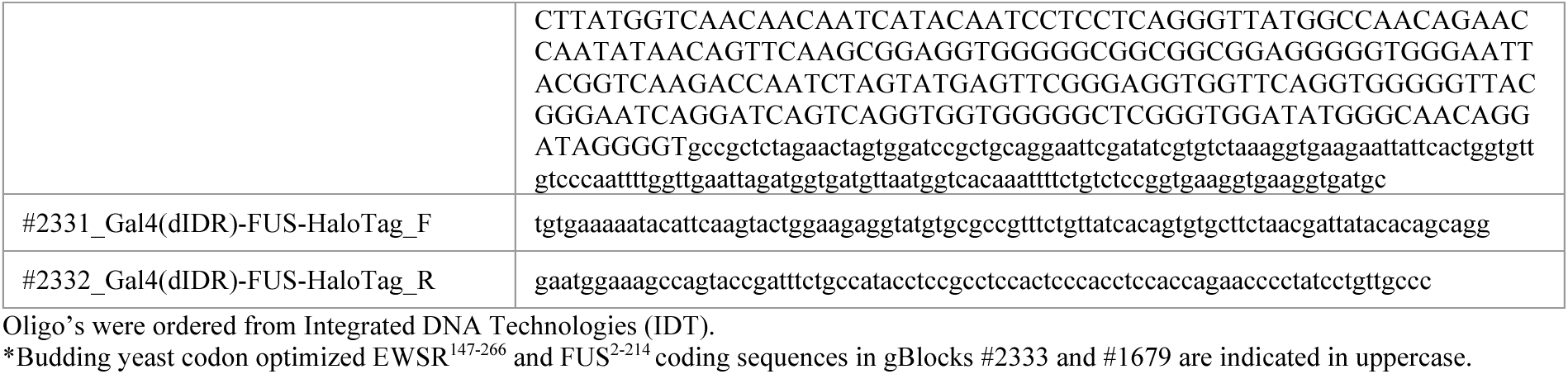
Oligos used in this study.

**Table S4.** Search time parameter estimates.

| | $t_{\text{search}}$<br>short (s)<br>± 95% CI | $t_{\text{search}}$<br>long (s)<br>± 95% CI | Fraction short<br>± 95% CI | Fraction<br>long<br>± 95% CI | Average<br>$t_{\text{search}}$ (s)<br>± 95% CI |
| --- | --- | --- | --- | --- | --- |
| Gal4 at *GAL* | 33.53 ± 8.41 | 557.51 ± 12.18 | 0.44 ± 0.03 | 0.56 ± 0.03 | 325.19 ± 18.58 |
| Gal4 at *RNR2* | 227.78 ± 60.95 | 3925.14 ± 1864.36 | 0.87 ± 0.07 | 0.13 ± 0.07 | 703.98 ± 350.41 |
| 1+1TFBS | 149.64 ± 15.87 | 787.53 ± 43.23 | 0.68 ± 0.03 | 0.32 ± 0.03 | 354.59 ± 23.88 |
| 1TFBS | 133.7 ± 11.88 | 1203.37 ± 136.46 | 0.86 ± 0.02 | 0.14 ± 0.02 | 280.87 ± 31.02 |
| 1TFBS+*tetO* | 57.0 ± 7.69 | 742.91 ± 45.22 | 0.77 ± 0.02 | 0.23 ± 0.02 | 218.05 ± 17.85 |
| Gal4 decreased | 79.35 ± 11.02 | 2175.19 ± 214.61 | 0.83 ± 0.02 | 0.17 ± 0.02 | 428.87 ± 53.07 |
| Gal4-DBD-only | 93.51 ± 16.62 | 2807.47 ± 427.22 | 0.83 ± 0.03 | 0.17 ± 0.03 | 543.15 ± 104.87 |
| Gal4ΔminiAD | 104.34 ± 7.81 | 1165.33 ± 88.47 | 0.86 ± 0.01 | 0.14 ± 0.01 | 257.13 ± 21.26 |
| Gal4ΔCR | 71.65 ± 10.74 | 827.0 ± 38.48 | 0.64 ± 0.02 | 0.36 ± 0.02 | 341.69 ± 22.18 |
| Gal4ΔIDR | 145.48 ± 11.68 | 2394.22 ± 213.66 | 0.84 ± 0.02 | 0.16 ± 0.02 | 494.36 ± 49.52 |
| Gal4ΔIDR-EWS | 90.76 ± 9.81 | 989.47 ± 70.82 | 0.8 ± 0.02 | 0.2 ± 0.02 | 274.52 ± 23.56 |
| Gal4ΔIDR-FUS | 91.53 ± 9.27 | 1000.64 ± 57.09 | 0.77 ± 0.02 | 0.23 ± 0.02 | 298.16 ± 21.47 |

